# Novel reporter of the PINK1-Parkin mitophagy pathway identifies its damage sensor in the import gate

**DOI:** 10.1101/2025.02.19.639160

**Authors:** Julia A. Thayer, Jennifer D. Petersen, Xiaoping Huang, James Hawrot, Daniel M. Ramos, Shiori Sekine, Yan Li, Michael E. Ward, Derek P. Narendra

## Abstract

Damaged mitochondria can be cleared from the cell by mitophagy, using a pathway formed by the recessive Parkinson’s disease genes PINK1 and Parkin. How mitochondrial damage is sensed by the PINK1-Parkin pathway, however, remains uncertain. Here, using a Parkin substrate-based reporter in genome-wide screens, we identified that diverse forms of mitochondrial damage converge on loss of mitochondrial membrane potential (MMP) to activate PINK1. Consistently, the MMP but not the presequence translocase-associated motor (PAM) import motor provided the essential driving force for endogenous PINK1 import through the inner membrane translocase TIM23. In the absence of TIM23, PINK1 arrested in the translocase of the outer membrane (TOM) during import. The energy-state outside of the mitochondria further modulated the pathway by controlling the rate of new PINK1 synthesis. Our results identify separation of PINK1 from TOM by the MMP, as the key damage-sensing switch in the PINK1-Parkin mitophagy pathway.

**Highlights:**

- MFN2-Halo is a quantitative single-cell reporter of PINK1-Parkin activation.
- Diverse forms of mitochondrial damage, identified in whole-genome screens, activate the PINK1-Parkin pathway by disrupting the mitochondrial membrane potential (MMP).
- The primary driving force for endogenous PINK1 import through the TIM23 translocase is the MMP with the PAM import motor playing a supporting role.
- Loss of TIM23 is sufficient to stabilize PINK1 in the TOM complex and activate Parkin.

## Introduction

Mitochondria use oxidative phosphorylation (OXPHOS) to generate most of the cell’s energy but suffer oxidative damage in the process ^1^. These damaged mitochondria accumulate with age, especially in long-lived neurons and myocytes – unless they can be recognized and degraded through a quality control process, such as the selective form of autophagy called “mitophagy” ^2^. One major mitophagy pathway eliminating damaged mitochondria is formed by the Parkinson’s disease (PD) genes PINK1 and Parkin ^3^. A robust understanding of how the PINK1-Parkin pathway is regulated may spur the development of new therapies for neurodegenerative disorders and other diseases resulting from mitochondrial damage.

In the PINK1-Parkin pathway, damaged mitochondria are first recognized by the ubiquitin kinase PINK1, using an elegant mechanism that relies on differential sorting of PINK1 by healthy and damaged mitochondria (reviewed in ^3^). Healthy mitochondria rapidly cleave PINK1 from their surface, by importing PINK1 along the TOM-TIM23 mitochondrial precursor path to the inner mitochondrial membrane (IMM). There, PINK1 is cleaved by the protease PARL and released from healthy mitochondria into the cytosol for degradation by the proteasome. Damaged mitochondria, by contrast, cannot fully import and cleave PINK1. Import arrested PINK1, instead, matures on the surface of the damaged mitochondria in a supercomplex with the outer membrane translocase TOM and inner membrane translocase TIM23 ^4–6^. There, PINK1 activates the E3 ubiquitin ligase Parkin in two steps.

PINK1, first, phosphorylates ubiquitin on outer mitochondrial membrane (OMM) proteins, which Parkin binds, and PINK1, then, directly phosphorylates Parkin on its ubiquitin like domain (UBL). Once activated, Parkin ubiquitinates several proteins on the OMM of the damaged mitochondrion. These, in turn, bind ubiquitin-dependent selective autophagy adaptors, including Optineurin ^7,8^, to initiate mitophagy. Some of Parkin’s most efficient substrates, including the mitochondrial fusion protein Mitofusin-2 (MFN2), can also be degraded by proteasome in the cytosol following their extraction from the OMM by AAA+-ATPase VCP and its adaptors ^9,10^.

Although many details of the PINK1-Parkin pathway have been worked out, several key questions remain, including, critically, how the PINK1-Parkin response is shaped by individual components of the TOM-TIM23 import precursor pathway and the two driving forces for mitochondrial protein import: the mitochondrial membrane potential (MMP) and (for some but not all precursors) the ATP-dependent presequence translocase-associated motor (PAM) complex ^11^. These questions are central to understanding how the PINK1-Parkin pathway senses mitochondrial damage. To help address these questions, we developed a novel screening approach for genome-wide activators and facilitators of the PINK1-Parkin pathway. We endogenously tagged one of Parkin’s preferred substrates, MFN2, with HaloTag ^12^, allowing us to monitor its degradation following PINK1-Parkin activation at the single cell level by flow cytometry. Notably, this strategy is complementary to existing single-cell reporters, such as the widely-used mitophagy reporters mt-Keima and mito-QC ^13,14^, allowing us to probe for PINK1-Parkin activators and facilitators that may have been missed by prior genome-wide screens.

Using this reporter in fluorescence-activated cell sorting (FACS)-based whole-genome CRISPRi screens (including in cells with endogenous Parkin for the first time), we uncovered several novel facilitators and activators of the PINK1-Parkin pathway. Together these suggest a model in which PINK1- Parkin activation is primarily sensitive to loss of the MMP, with the ATP-dependent PAM complex playing a supporting role. The MMP, in this model, provides the main driving force allowing the TIM23 translocase to separate PINK1 from its stable interaction with TOM. We propose this mechanism is the “damage sensor” in the PINK1-Parkin pathway.

## Results

### MFN2-Halo is a sensitive single-cell reporter of PINK1-Parkin activation

As PINK1-Parkin activation robustly degrades MFN2 upon activation, we reasoned that whole-cell MFN2 levels might be used as a sensitive single-cell reporter of PINK1-Parkin activity (Fig. 1A). To test this idea, we endogenously tagged MFN2 with HaloTag (MFN2-Halo), in a HeLa cell line that additionally expressed dCas9-BFP-KRAB or dCas9-BFP-ZIM3 to enable CRISPR interference (CRISPRi) ^15,16^, with or without exogenous mCherry-Parkin (mCh-Parkin).

**Figure 1.**
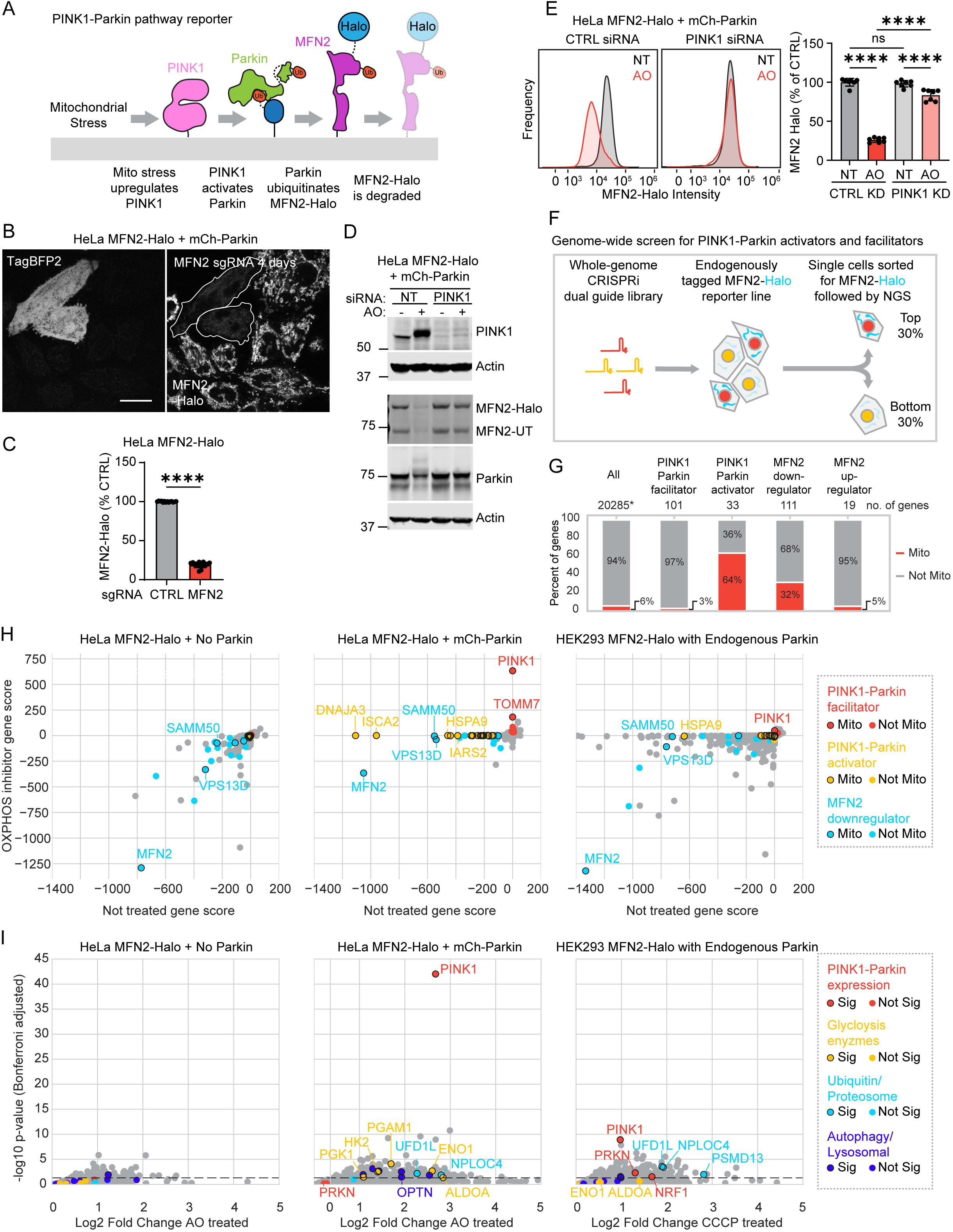
Genome-wide screens with the MFN2-Halo reporter identify activators and facilitators of the PINK1-Parkin mitophagy pathway. (A) Schematic illustrating the PINK1-Parkin pathway reporter MFN2-Halo. (B) Representative confocal images of HeLaMFN2-Halo+mCh-Parkin cells transduced with a cassette co-expressing a dual sgRNA targeting MFN2 and TagBFP2 to mark transduced cells (white outline). Scale bar = 20 µm. (C) Flow cytometry of HeLaMFN2-Halo cells. **** p ≤ 0.0001 Error bars mean +/- SD. N = 3 independent experiments. (D) Representative immunoblots of endogenously tagged and untagged MFN2 alleles (MFN2-Halo and MFN2-UT, respectively) of same cells in (1B). N = 5 replicates on at least 2 occasions. (E) Flow cytometry of HeLaMFN2-Halo+mCh-Parkin cells +/- AO treatment. **** p ≤ 0.0001 Error bars mean +/- SD. N = 3 independent experiments. (F) Schematic illustrating the FACS based genome-wide CRISPRi screening strategy. Six screens were performed in total, using three different cell lines with or without exposure to OXPHOS inhibitors. (G) Stacked bar graph represents the proportion of nuclear gene perturbations encoding mitochondrial (mito) vs. non-mitochondrial proteins (not mito). (H) Dot plots of gene scores (product of log2 fold change and adjusted -log10 p-value) from screens described in (F). (I) One-sided volcano plots for screens described in (F).

After incubation with the Halo ligand Janelia Fluor 646, HeLa^dCas9-BFP-KRAB+MFN2-Halo+mCherry-Parkin^ (henceforth HeLa^MFN2-Halo+mCh-Parkin^) cells exhibited fluorescence in an OMM pattern consistent with MFN2’s known localization ^17^ (Fig. 1B). This pattern was absent in cells expressing a guide RNA (sgRNA) targeting endogenous MFN2 (Fig. 1B). Similarly, expression of a MFN2 sgRNA in HeLa^dCas9-BFP-KRAB+MFN2-Halo^ cells (henceforth HeLa^MFN2-Halo^) caused a ∼5-fold decrease in MFN2-Halo fluorescence measured at the whole cell level by flow cytometry (Fig. 1C), indicating that MFN2 protein levels could be determined in single cells over a wide dynamic range. Similar results were obtained in cells fixed immediately after CCCP treatment and stored for a day at 4°C prior to sorting (Fig. S1A). Immunoblotting for MFN2 showed the tagged and untagged copies of MFN2 were degraded by Parkin to a similar extent following Parkin activation by OXPHOS inhibition with antimycin and oligomycin (AO) (Fig. 1D). MFN2-Halo degradation by immunoblotting and flow cytometry was blocked by PINK1 knockdown (KD), demonstrating that it is mediated by the PINK1-Parkin pathway in HeLa cells (Fig. 1D and E). We then endogenously tagged MFN2 with HaloTag in HEK293^dCas9-BFP-ZIM3^ cells (henceforth HEK293^MFN2-Halo^), which express endogenous Parkin at low levels (nTPM = 0.5, Human Protein Atlas proteinatlas.org). As expected, PINK1-dependent degradation of MFN2-Halo was also detected in HEK293^MFN2-Halo^ cells treated with 10 µM CCCP, albeit at a lower level and with slower kinetics than in HeLa^MFN2-Halo+mCh-Parkin^ cells (Supplemental Fig. S1B - C).

### Genome-wide CRISPRi screens with MFN2-Halo reporter identifies facilitators and activators of the PINK1-Parkin pathway

Next, we screened for PINK1-Parkin activators and facilitators in a pooled FACS-based format. The three MFN2-Halo reporter cell lines were transduced with a whole-genome dual guide CRISPRi library and left untreated or treated with OXPHOS inhibitors (AO or CCCP) (Fig. 1F).

Comparison of HeLa^MFN2-Halo^ screens with and without mCh-Parkin distinguished PINK1-Parkin activators and facilitators from guides that alter MFN2-Halo levels independently of the PINK1-Parkin pathway (Fig. 1G – I and Table S1). Activators were those perturbations that caused Parkin-dependent MFN2-Halo degradation in the untreated condition; facilitators were those perturbations that blocked Parkin-dependent MFN2-Halo degradation following OXPHOS inhibition. The top facilitators included genes known to be critical for activation of the PINK1-Parkin pathway, including PINK1; TOMM7, which stabilizes PINK1 on OMM ^18^; and OPTN and the VCP adaptors NPLOC4 and UFD1L, which degrade MFN2 following PINK1-Parkin activation ^7–10^. Strikingly, most PINK1-Parkin activators encoded mitochondrial proteins, and many have not been previously reported as PINK1-Parkin activators (Fig. 1G). These included, surprisingly, TIMM23, a core subunit of the TIM23 complex, as explored further below.

Overall, the gene scores were substantially higher in the HeLa^mCh-Parkin^ screens than the HEK293 screens with endogenous Parkin. This was expected given the more robust degradation of MFN2-Halo in the presence of high Parkin levels. Conversely, the HEK293 screen revealed genes, such as Parkin and NRF1, which are required to maintain endogenous Parkin expression ^19^. These were absent from the HeLa screen, in which Parkin was expressed from an exogenous promoter.

Together these datasets provide a deep resource of genes modulating the PINK1-Parkin pathway, including (for the first time) the endogenous PINK1-Parkin pathway. They additionally provide a valuable resource for genome-wide regulators of MFN2 expression, independent of the PINK1-Parkin pathway. Among the notable mitochondrial pathways downregulating MFN2 expression in both HeLa and HEK293 cells were mitochondrial fission (DNM1L and MFF), phospholipid transfer and metabolism (VPS13D, TAZ1), PINK1-Parkin independent mitophagy (FBXL4, TMEM11), and cristae remodeling (SAMM50, MINOS, IMMT, ATP5H, ATP5O, ATPC1, ATP5L, ATP5I, ATP5A1, ATP5D) (Table S1).

### Glycolysis facilitates activation of the PINK1-Parkin pathway following OXPHOS impairment

We focused our detailed investigations initially on facilitators of the PINK1-Parkin pathway. Notable among these were genes encoding five core glycolytic enzymes (ENO1, ALDOA, HK2, PGK1, and PGAM1), some of which were also identified in the endogenous screen (but did not reach genome-wide significance) (Fig. 1I). Although HK2 has previously reported as a PINK1 facilitator ^20,21^, with a proposed mechanism that is independent of its role in glycolysis ^21^, our identifying four other glycolytic enzymes suggested that glycolysis may serve a broader function in PINK1-Parkin activation than previously recognized.

To explore this, we focused first on ENO1 which had the strongest effect in the screen and catalyzes the penultimate reaction in the canonical glycolysis pathway (Fig. 2A). ENO1 KD blocked Parkin dependent degradation of MFN2 in HeLa^MFN2-Halo+mCh-Parkin^ to a similar extent as PINK1 KD in HeLa^MFN2-Halo+mCh-Parkin^ cells treated with the mitochondrial uncoupler CCCP (Fig., 2B, top). Similarly, ENO1 KD blocked PINK1-Parkin mitophagy in HeLa^dCas9-BFP-KRAB+mt-Keima^ (hereafter HeLa^mt-Keima^) cells treated with CCCP (Fig. 2B, middle). Together, this confirmed that glycolytic enzymes are required for mitochondrial ubiquitination and mitophagy by the PINK1-Parkin pathway following OXPHOS inhibition.

**Figure 2.**
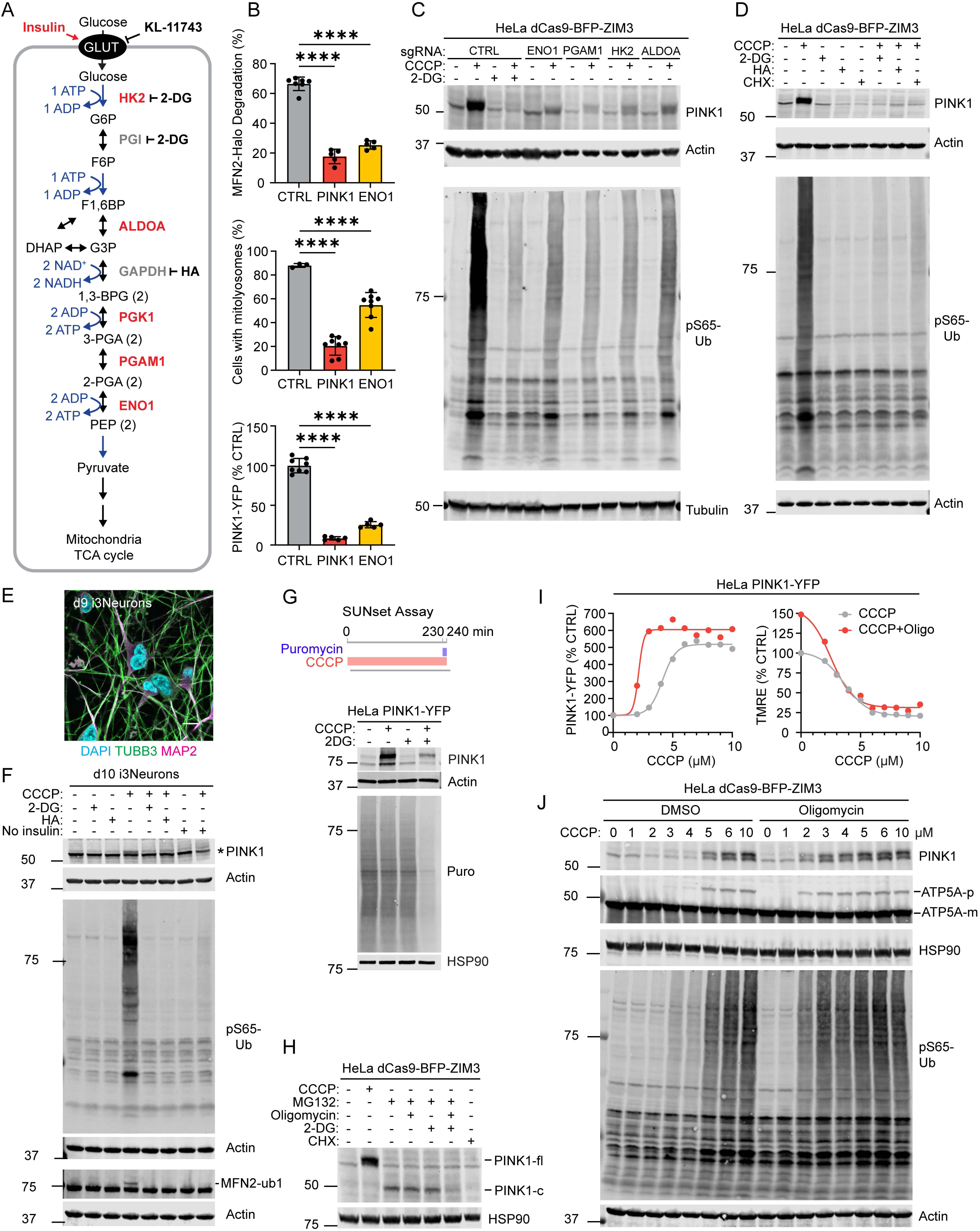
Glycolysis is required for new PINK1 translation and activation following OXPHOS deficiency. (A) Schematic illustrating the core glycolysis pathway with screen hits in red and inhibitors tested in black. (B) Flow cytometry measurements in HeLa cells treated with CCCP 10 µM for 4 hrs. **** p ≤ 0.0001 Error bars mean +/- SD. N = 6 independent experiments from two separate transductions. (C) Representative immunoblots of HeLadCas9-BFP-ZIM3 cells treated with 10 µM CCCP +/- 10 mM 2-DG for 4 hrs. N = 3 independent experiments. (D) Representative immunoblots of HeLadCas9-BFP-ZIM3 cells treated with 10 µM CCCP, 10 µM HA, 10 mM 2- DG, and/or 50 µg/mL CHX for 4 hrs. N = 3 independent experiments. (E) Representative confocal image. Scale bar = 10 µm. (F) Representative immunoblot of lysates from i3Neurons, treated with 10 mM 2-DG, 10 µM HA, and/or 20 µM CCCP, with or without of insulin withheld. * denotes non-specific band. MFN2-ub1 = monoubiquitinated MFN2. N ≥ 6 total replicates from at least 2 independent differentiations. (G) Immunoblot SUNset assay (bottom) performed in HeLaPINK-YFP cells as indicated in scheme (top). Puro = puromycylated proteins. N = 3 independent experiments. (H) Immunoblot of cleaved PINK1 (PINK1-c) abundance, stabilized by proteasome inhibition (50 µM MG132), following inhibition of mitochondrial ATP (10 µg/mL oligomycin) and/or glycolytic ATP (10 mM 2-DG) production for 4 hrs. N = 3 replicates on at least 2 occasions. (I) Flow cytometry measurements of PINK1-YFP (left graph) and mitochondrial membrane potential with TMRE 20 nM (right graph), following drug treatment for 4 hrs. Graphs are representative of N = 3 independent experiments. (J) Immunoblot of HeLadCas9-BFP-ZIM3 cells treated with escalating doses of CCCP with or without F1FO-ATP synthase inhibition (oligomycin 10 µg/mL) for four hours. Import block of ATP5A is monitored through the accumulation of uncleaved ATP5A (ATP5A-p) relative to mature ATP5A (ATP5A-m). N = 3 independent experiments.

We next assessed whether ENO1 has its effect up- or down-stream of PINK1, by monitoring PINK1-YFP levels in HeLa^PINK1KO+PINK1-YFP+dCas9-BFP-ZIM3^ (henceforth HeLa^PINK1-YFP^) cells. ENO1 KD dramatically blocked the increase in PINK1-YFP following CCCP treatment (Fig. 2B, bottom). This demonstrated that

ENO1 is required at the first step of the PINK1-Parkin pathway, in which newly translated PINK1 is stabilized on the OMM ^22^. PINK1-YFP accumulation following OXPHOS inhibition with either CCCP, AO, or rotenone (a complex I inhibitor) + oligomycin (RO) was also blocked by the HK2 inhibitor 2- deoxyglucose (2-DG) (Fig. S1D).

We next tested the effect of the four strongest glycolysis hits (ENO1, PGAM1, HK2, and ALDOA) on stabilization of endogenous PINK1 on uncoupled mitochondria (Fig. 2C). Knockdown of each blocked accumulation of endogenous PINK1 and PINK1’s substrate phospho-S65 ubiquitin (pS65-Ub). Inhibition of HK2 with 2-DG or GAPDH with heptelidic acid (HA) similarly blocked accumulation of endogenous full-length PINK1 in HeLa cells (Fig. 2D) and in neuron-like cells induced from iPSCs by neurogenin-2 expression (i^3^Neurons) ^23^ (Fig. 2E and F). Blocks upstream of glycolysis (glucose starvation and inhibition of glucose uptake) phenocopied inhibition of glycolysis (Fig. S1E and F), as did withdrawal of insulin, which maintains neuronal glucose uptake ^24^, in i^3^Neurons (Fig. 2F, lane 8 vs. 4). Bypassing glycolytic ATP production with galactose, which is oxidized to pyruvate without net ATP production, or supplementing glucose-free media with high pyruvate did not rescue the PINK1-YFP response to OXPHOS inhibition (Fig. S1F). This points to glycolytic ATP production as the critical requirement for PINK1 accumulation. In contrast to glycolysis inhibitors, SYNJ2, recently implicated in PINK1 mRNA transport ^25,26^, did not block PINK1 synthesis in HeLa cells (Fig. S1G). HK2 and ENO1 KD decreased PINK1-YFP incorporation into the TOM complex, measured by clear native PAGE (CN-PAGE), in proportion with their effects on total PINK1 accumulation (Fig. 2C compared to Fig. S1H), arguing against a selective effect of HK2 on incorporation of PINK1-YFP into the TOM complex, as previously proposed ^21^. Together these results demonstrate that glycolysis is required for PINK1 accumulation and activation following OXPHOS inhibition in HeLa cells and human i^3^Neurons, likely through the glycolytic production of ATP.

We next considered why glycolytic ATP is required for PINK1 accumulation following OXPHOS inhibition. Activation of the PINK1-Parkin pathway requires new PINK1 to be synthesized for accumulation on impaired mitochondria and can be blocked by the ribosome inhibitor cycloheximide (CHX) ^22^ (Fig. 2D, lane 8 vs. 2). As protein synthesis is energy demanding ^27^, we hypothesized that glycolysis may be needed in the setting of OXPHOS inhibition to provide energy for new PINK1 synthesis. To test this hypothesis, we monitored global protein translation in the setting of glycolysis and OXPHOS inhibition, using the SUNset assay ^28^. Consistent with high energy requirements for translation, 10 mM 2-DG + 10 µM CCCP inhibited global protein translation in parallel with the block in full length PINK1 accumulation (Fig. 2G, lane 4 vs. 2 and Fig. S1I). To test this another way, we monitored the accumulation of PARL-cleaved PINK1 ^29^, stabilized by the proteosome inhibitor MG132. Inhibition of both mitochondrial ATP production (using the F_1_F_O_-ATP synthase inhibitor oligomycin) and glycolytic ATP production with 2-DG dramatically decreased production of cleaved PINK1 (Fig. 2H, lane 6 vs. 3).

Together these findings demonstrate that glycolytic ATP production is required for new PINK1 synthesis in the setting of reduced OXPHOS to activate the PINK1-Parkin pathway.

The mitochondrial ATP pool is partially separate from the cytosolic ATP pool and therefore may have distinct effects on PINK1-YFP stabilization in response to MMP lowering with CCCP. To test these effects, we inhibited mitochondrial ATP production with oligomycin (a selective inhibitor of the F_1_F_O_-ATP synthase) in conjunction with increasing concentrations of CCCP, while simultaneously monitoring MMP and PINK1-YFP intensity by flow cytometry. Notably, oligomycin was not sufficient to stabilize PINK1-YFP on its own but lowered the MMP threshold for PINK1-YFP stabilization (Fig. 2I). As mitochondrial ATP contributes to import along the precursor path through cycling of HSPA9 (also known as, mt-HSP70) in the PAM import motor ^11^, we hypothesized that oligomycin may sensitize mitochondria to MMP treatment by removing the other driving force, the PAM import motor, for precursor import. Consistent with this hypothesis, oligomycin co-treatment lowered the threshold for import block of both endogenous ATP5A and endogenous PINK1 by CCCP (Fig. 2J).

Together these results demonstrate that the overall energetic state of the cell modulates PINK1- Parkin activation through its effects on PINK1 synthesis: ATP generated by glycolysis becomes limiting for new PINK1 synthesis in the setting OXPHOS dysfunction, attenuating the PINK1-Parkin response.

Selectively lowering mitochondrial ATP, by contrast, reduces the MMP threshold for the PINK1-Parkin pathway activation.

### Diverse forms of mitochondrial damage activate the PINK1-Parkin pathway by disrupting the MMP

We turned next to the PINK1-Parkin activators identified in the screen (Fig. 3A). To identify points of convergence among the functionally diverse activators we compared their impact on mitochondrial ultrastructure, the mitochondrial proteome, quantitative PINK1-Parkin activity reporters, and the MMP (Fig. 3B).

**Figure 3.**
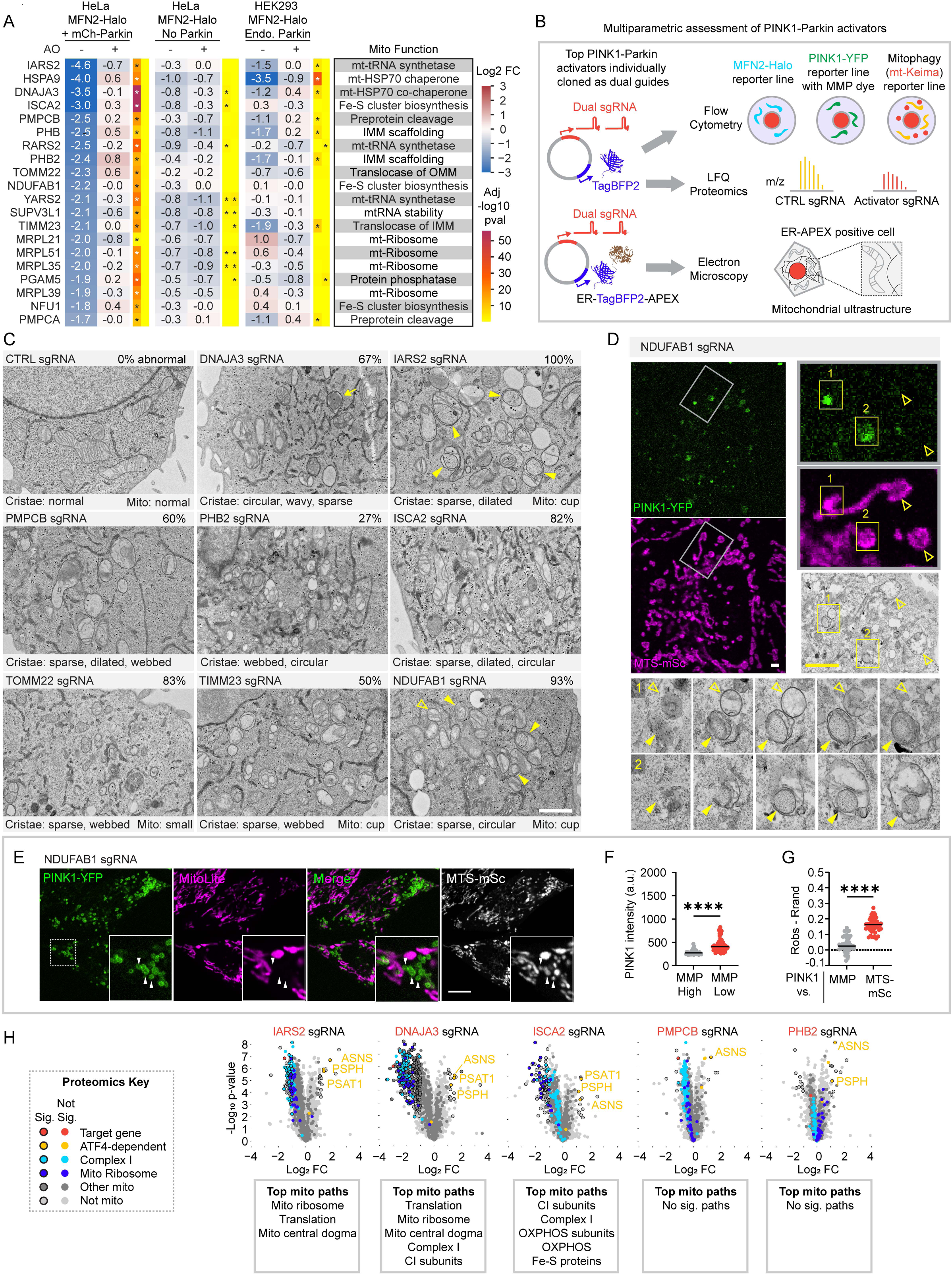
PINK1-Parkin activators reflecting diverse mitochondrial functions converge on damage to the inner mitochondrial membrane. (A) Heat maps show log2 fold changes for the top 20 mitochondrial PINK1-Parkin activators in each of the six screens described in (Fig. 1F). (B) Schematic summarizing the methods used to characterize the top PINK1-Parkin activators. (C) Representative TEM micrographs show abnormalities in mitochondrial ultrastructure induced by knockdown of the top PINK1-Parkin activators in HeLadCas9-BFP-ZIM3 cells transduced with sgRNA and APEX- ER. Transduced cells were readily identified by dark staining of the osmophilic polymer in the ER lumen produced by the APEX-DAB reaction. In some samples, cupped mitochondria (closed yellow arrowheads), indicative of membrane potential collapse 31,32, were found adjacent to intact mitochondria (open arrowheads). A fluffy aggregate in the matrix of mitochondria (arrow) was observed in DNAJA3 sgRNA cells. The percentage of cells with abnormal mitochondria out ten or more cells imaged per sample is indicated in the upper right of each image. Text below images describes the predominant abnormal features of the cristae and cup-shaped or small mitochondrial morphology if observed. Control cells from two transductions were analyzed, while single transductions were performed and analyzed for non-control cells. Scale bar = 1 µm. (D) CLEM images from a NDUFAB1 KD HeLaPINK1-YFP+MTS-mSc cell. PINK1-YFP selectively accumulates on cupped mitochondria (closed arrowheads) while sparing adjacent intact mitochondria (open arrowheads). Serial sections of two boxed areas (1 and 2) are shown in magnifications. Scale bars = 2 µm. (E) Representative confocal image of live HeLaPINK1-YFP+MTS-mSc cells with NDUFAB1 KD, showing PINK1-YFP accumulation on a subset of mitochondria that have not accumulated the MMP-dependent dye MitoLite NIR (arrow heads). Scale bar = 10 µm. (F) Graph of cells in (E) comparing PINK1-YFP intensity on mitochondria with and without MMP within the same cell, measured from confocal images as in (E). **** p ≤ 0.0001. N = 48 cells were analyzed in total from 5 wells and 2 independent transductions. (G) Graph of cells in (E) comparing PINK1-YFP co-localization to MTS-mSc and the MMP dye MitoLite NIR. The mean Pearson coefficient for 20 pixel-randomized images (Rrand) was subtracted from the measured Pearson coefficient (Robs) for each image. **** p ≤ 0.0001. N = 50 cells were analyzed in total from 5 wells and 2 independent transductions. (H) Volcano plots show changes to whole cell protein abundance following transduction with the indicated guide vs. a non-targeting guide in HeLa cells without Parkin (HeLadCas9-BFP-ZIM3). Boxes show top MitoCarta3.0 mitochondrial pathways identified in enrichment analysis of significantly down-regulated proteins. N = 4 replicates/sgRNA on 1 occasion.

We first considered their effects on mitochondrial ultrastructure by transmission electron microscopy (TEM), using a dual guide vector we modified to co-express ER targeted APEX, a genetically encoded reporter for TEM ^30^ (Fig. 3B, bottom). Transduced HeLa^dCas9-BFP-ZIM3^ cells could be readily identified at the TEM level by the darkly stained osmophilic polymer in the ER lumen produced by the APEX-DAB reaction (Fig. 3C and Fig. S2A and B). Comparison of mitochondrial ultrastructure among the top activators demonstrated diverse forms of mitochondrial damage that reflected the known mitochondrial function of the targeted genes—e.g., matrix protein inclusions following loss of the matrix mt-HSP70 co-chaperone DNAJA3 and disruption of the boundary IMM following loss of the scaffolding protein PHB2 (Fig. 3C, Fig. S2C, and Table S2). However, they shared disruption of mitochondrial cristae – the enfolded region of the IMM that houses the OXPHOS machinery – in at least a portion of the mitochondrial network. In control cells, cristae were studded with protein densities that are likely the F_1_F_O_-ATP synthase; these protein densities were rare or absent on the disrupted cristae (Fig. S3). A subset of mitochondria in several of the knockdowns (including NDUFAB1 and IARS2) formed abundant ring or cup shapes (hereafter, cupped mitochondria) (Fig. 3C, yellow closed arrowheads in upper right and lower right images, and Fig. S2C), phenocopying severe loss of membrane potential following treatment with CCCP ^31,32^. These cupped mitochondria were often adjacent to intact mitochondria (Fig. 3C, yellow open arrowheads), reflecting a stochastic mechanism of mitochondrial damage or the ability to concentrate limited working OXPHOS components into a subset of functional mitochondria. Using correlative light electron microscopy (CLEM), PINK1-YFP was found to accumulate selectively on these cupped mitochondria following knockdown of NDFUAB1 in HeLa^PINK1-YFP+MTS-mScarlet^ cells (henceforth HeLa^PINK1-YFP+MTS-mSc^) (Fig. 3D). Consistent with these representing depolarized mitochondria, analysis of live cells demonstrated that the low MMP mitochondrial population had higher PINK1-YFP than the high MMP mitochondrial population in the same cell (Fig. 3E and F). Similarly, PINK1-YFP correlated with MTS-mSc but poorly with MMP (Fig. 3G). Considered together, the mitochondrial ultrastructure was diverse among the activators, but shared disruption of the mitochondrial cristae, the MMP-generating structure of mitochondria, and morphologies characteristic of severe loss of MMP, such as cupping; PINK1 preferentially targeted the de-energized, cupped mitochondria in cells with a heterogenous mitochondrial population.

Disruptions to the proteomes of five activators tested were similarly varied and reflected the known function of the targeted genes (Fig. 3H and Table S3). Knockdown of IARS2, a mitochondrial tRNA synthetase, caused a severe loss of proteins related to mitochondrial translation, including those comprising the mitochondrial ribosome; loss of ISCA2 destabilized complex I subunits and other proteins containing Fe-S clusters; disrupting the matrix co-chaperone DNAJA3 destabilized matrix and IMM proteins *en mass*, while PMPCB and PHB2 knockdowns were minimally disruptive to mitochondrial protein abundance. In contrast to these diverse impacts on the mitochondrial proteome, all five PINK1- Parkin activators caused a global decrease in mitochondrial proteins, when comparing HeLa cells expressing and not expressing mCh-Parkin, likely due to increased PINK1-Parkin mitophagy (Fig. S3B). Additionally, all five activated the OMA1-DELE1-ATF4 integrated stress response (mt-ISR), evidenced by increased expression of ATF4 target gene products ASNS, PSAT1, and/or PSPH ^33–35^ (Fig. 3H). This is consistent with stress sensed at the IMM by the OMA1-DELE1 pathway ^34,35^, in parallel with activation of the PINK1-Parkin mitophagy pathway. Thus, the PINK1-Parkin activators, though varied in their perturbations to mitochondrial structure and protein composition, converged on stress to the IMM and activation of damage responses.

To further pinpoint the point of convergence among the PINK1-Parkin activators, we next assessed the impact of 18 of the 20 using a suite of quantitative flow cytometry-based reporters we enabled for CRISPRi (Fig. 3B, top scheme). Notably, knockdown of the top screen hits caused MFN2- degradation (HeLa^MFN-Halo+mCh-Parkin^), PINK1-YFP stabilization (HeLa^PINK1-YFP^), and loss of MMP in a correlated and graded fashion, with knockdown of some targets (group 1) having little effect and others (group 4) having a strong effect across the three measures (Fig. 4A – C). Comparison of PINK1-YFP and MMP measured in the same experiments showed a non-linear inverse relationship between MMP and PINK1- YFP, with PINK1-YFP markedly increasing after MMP dye intensity was reduced to ∼50% of control levels (Fig. 4C). Strikingly, this followed a similar pattern as observed with increasing doses of the mitochondrial uncoupler CCCP (Fig. 2I). Notably, activators in groups 3 and 4 are involved in pathways critical for the maturation of newly imported matrix and IMM targeted proteins, including import (TOMM22 and TIMM23), folding of newly imported proteins (HSPA9 and DNAJA3), Fe-S cluster biosynthesis (ISCA2 and NDUFAB1), and cleavage of the preprotein sequence (PMPCA and PMPCB). This identifies maturation of newly important proteins as a critical vulnerability leading to MMP loss and PINK1-Parkin activation. Top activators were also found to promote PINK1-Parkin mitophagy, measured with the mt-Keima reporter (Fig. 4D), and to activate endogenous PINK1, measured by immunoblotting (Fig. S4A). Interestingly, although these mitochondrial perturbations strongly reduced MMP, they correlated only weakly with mitochondrial perturbations previously found to lower respiratory ATP production or increase mitochondrial superoxide production (e.g., genes encoding CI-, III-, and CoQ10 biosynthesis-related proteins), identified in recent systematic screens that utilized a similar CRISPRi library ^36,37^ (Fig. 4E and Table S4). Together these findings suggest a model in which functionally diverse sources of mitochondrial damages converge on disruption of MMP and subsequent stabilization of PINK1 as the primary mechanism of PINK1-Parkin activation.

**Figure 4.**
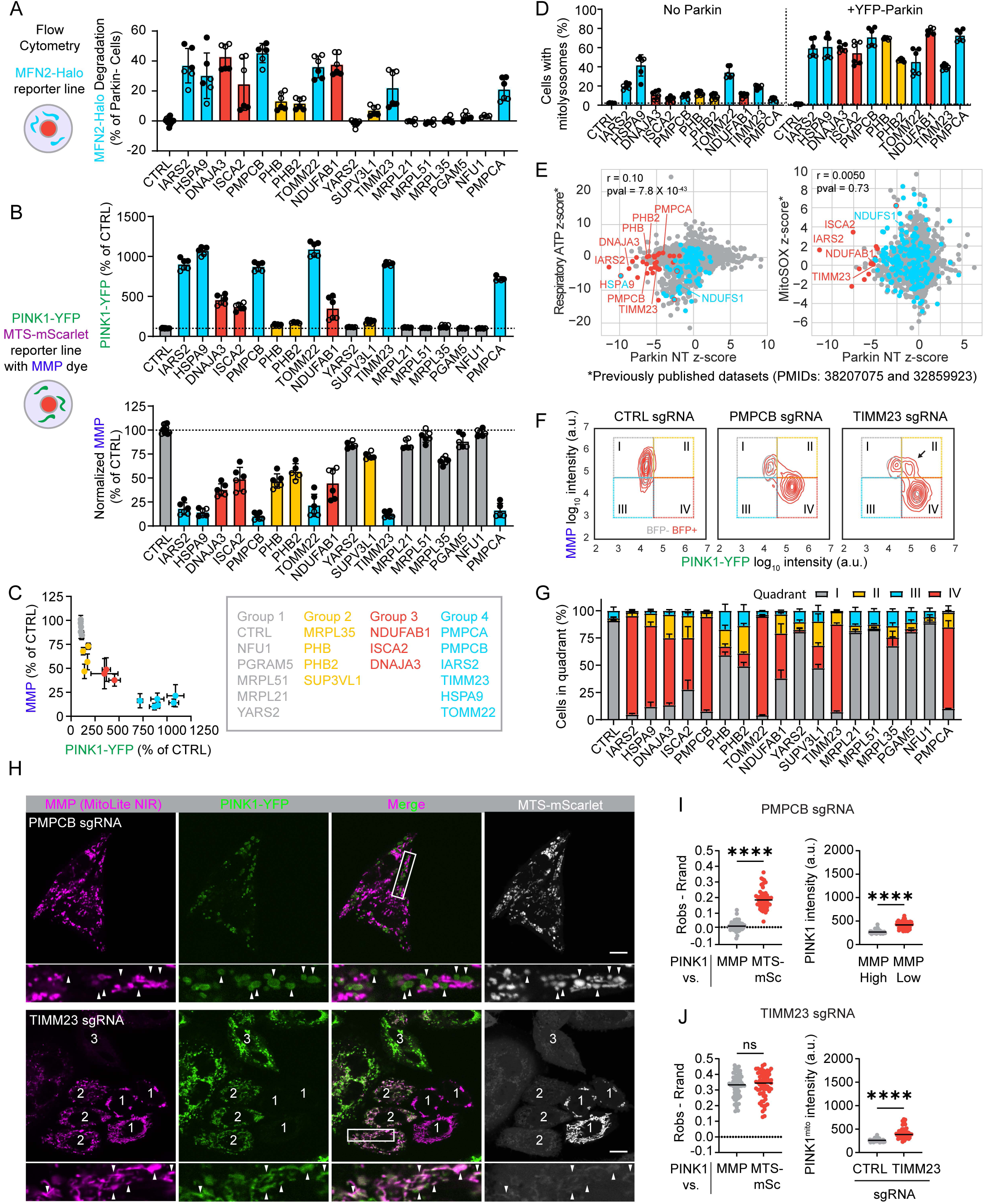
Diverse mitochondrial stresses disrupt the mitochondrial membrane potential to activate the PINK1-Parkin pathway. (A) Flow cytometry measurements of MFN2-Halo degradation (HeLaMFN2-Halo+mCh-Parkin cells). N = 6 biological replicates measured from 2 independent transductions. (B) Flow cytometry measurements of PINK1-YFP (top) and MMP with dye MitoLite NIR (bottom) obtained from same cells. (HeLaPINK1-YFP cells) N = 6 biological replicates measured from 2 independent transductions. (C) Graph of experiment in (B) directly comparing PINK1-YFP and MMP for each gene perturbation. (D) Flow cytometry measurements of mitophagy using HeLamt-Keima cells without Parkin (left) or with exogenous YFP-Parkin expression (right). N = 6 biological replicates measured from 2 independent transductions. (E) Dot plots comparing z-scores from the MFN2-Halo CRISPRi screen in not treated HeLa cells expressing mCh-Parkin (Fig. 1H, middle) with previously published CRISPRi screens of respiratory ATP (left) and mitochondrial superoxide (right) 36,37. (F) Representative 2D kernel density plots comparing single-cell PINK1-YFP intensity and intensity of the MMP sensitive dye MitoLite NIR from the same experiments as shown in (Fig. 4B and C). Arrow indicates a distinct population of cells, high in both MMP and PINK1-YFP, that was observed following TIMM23 KD. (G) Quantification of the proportion of cells in each of the quadrant 2D kernel density plots, such as those seen in (Fig. 3F and Fig. S4B). (H) Representative confocal images obtained as in (Fig. 3E). Scale bar = 10 µm. (I) Graph on (left) is as described in (Fig. 3G). **** p ≤ 0.0001. N = 54 cells were analyzed in total from 4 wells and 2 separate transductions. Graph on (right) is as described in (Fig. 3F). **** p ≤ 0.0001. N = 54 cells were analyzed in total from 4 wells and 2 independent transductions. (J) Graph on (left) is as described in (Fig. 3G). **** p ≤ 0.0001. N = 65 cells were analyzed in total from 6 wells and 2 separate transductions. Graph on (right) compares PINK1-YFP intensity on mitochondria following CTRL or TIMM23 KD. Only cells with high MMP were analyzed from both groups and only cells with mitochondrial block (based on MTS-mSc signal) were analyzed for the TIMM23 KD group. **** p ≤ 0.0001. N = 65 TIMM23 KD cells and 82 CTRL KD cells were analyzed in total from 6 wells and 2 independent transductions.

This conclusion was further supported by analysis of PINK1-YFP and MMP at the single cell level (Fig. 4F and G and Fig. S4B). Among the 18 activators, single cells were generally found in one of two states: high MMP / low PINK1-YFP or low MMP / high PINK1-YFP. Notably, this pattern was also observed with knockdown of the mitochondrial processing protease (MPP) subunits PMPCB and PMPCA, which were previously reported to stabilize PINK1 in the absence of MMP lowering ^38^. Live confocal microscopy confirmed that PINK1-YFP accumulated preferentially on the population of de-energized mitochondria within single cells following PMPCB knockdown (Fig 4H, top, and 4I), similar to the pattern observed for NDUFAB1 above. Thus, PINK1-YFP accumulation correlated closely with loss of the MMP for most PINK1-Parkin activators.

The pattern was different, however, for TIMM23 KD. Instead, a unique population of cells was observed with both high MMP and elevated PINK1-YFP (Fig. 4F, arrow). To explore this further, we examined TIMM23 KD in these cells by live confocal microscopy. Consistent with the flow cytometry results, a population of cells was observed with blocked import of MTS-mSc, high PINK1-YFP expression, and high MMP (Fig. 4H, bottom, cells numbered “2”, and 4J). PINK1-YFP expression was uniformly elevated throughout the mitochondrial network with retained MMP, reflected in the correlation between PINK1-YFP and MMP (Fig. 4J, left). This suggests a distinct progression following TIMM23 KD. PINK1-YFP initially accumulates on import blocked mitochondria that are still able to maintain MMP (Fig. 4H, bottom, cells numbered “2”). As the import block continues, however, OXPHOS fails, and the MMP is reduced (Fig. 4H, bottom, cell numbered “3”). This sequence differs from other activators, exemplified by NDUFAB1 and PMPCB, in which MMP loss precedes and likely causes import block.

Considered together, these findings demonstrate that import block through the TIM23 complex (either from TIMM23 KD or loss of the driving force for import through the TIM23 translocase) is the primary trigger for PINK1-YFP accumulation on mitochondria.

### PINK1 is stabilized in TOM complex following loss of TIM23 translocase

PINK1 was recently suggested to form a supercomplex with the TOM and TIM23 translocases following loss of the MMP ^4,5^, which was proposed to stabilize PINK1 on the OMM. It was, thus, surprising to observe that TIMM23 KD, itself, causes PINK1-YFP accumulation. To explore this further, we first verified that PINK1-YFP binds both the TOM and TIM23 translocases following loss of MMP, by affinity purification mass spectrometry (AP-MS). Consistent with PINK1-YFP forming a supercomplex with the TOM and TIM23 translocases, PINK1-YFP pulled down core components of the TOM complex (TOMM40 and TOMM22) and the TIM23 translocase (TIMM23 and TIMM17B) (Fig. 5A and Table S5).

**Figure 5.**
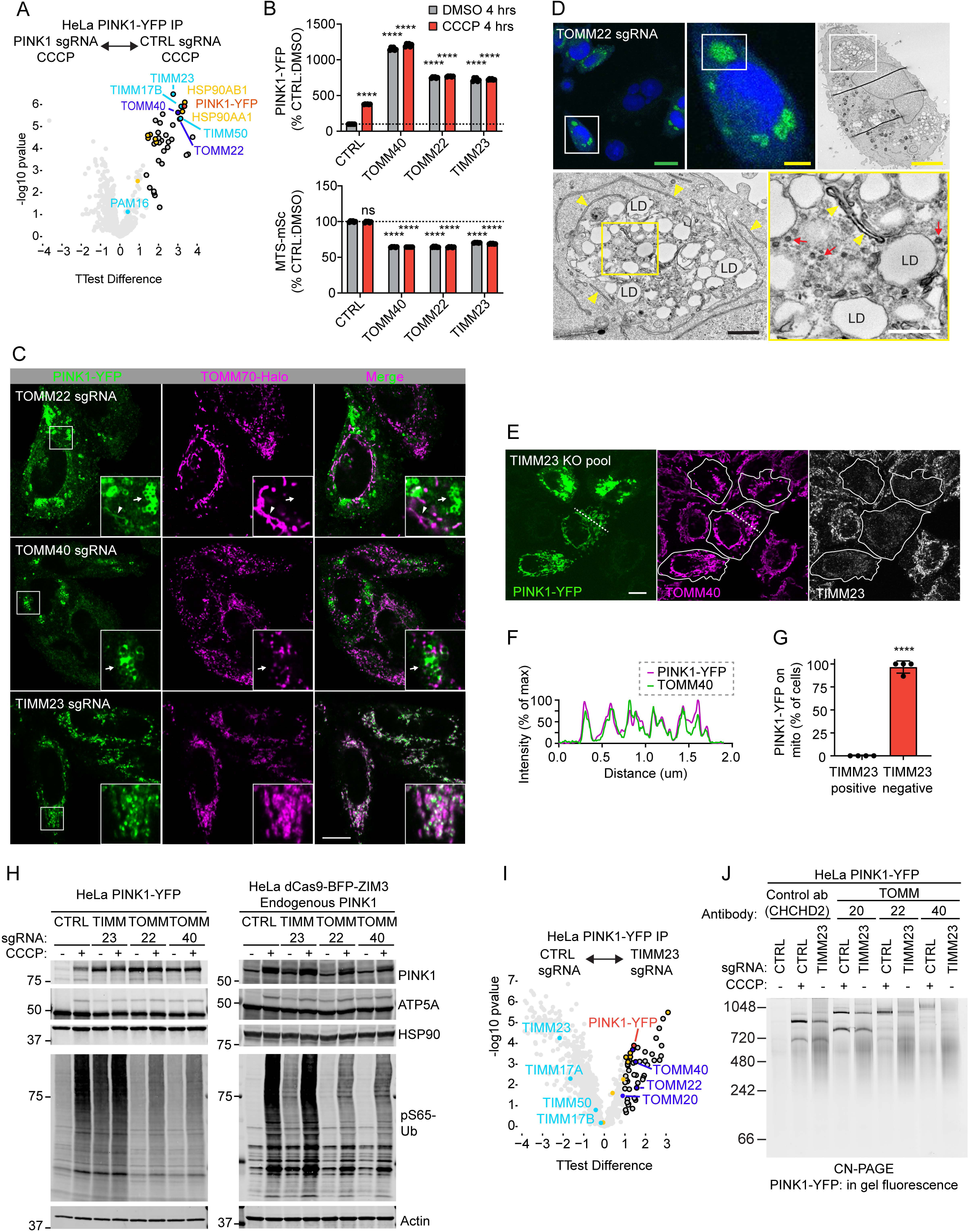
PINK1 stabilization on the OMM requires the TOM but not the TIM23 translocase. (A) Volcano plot shows interactors of PINK1-YFP (red) identified by AP-MS following transduction with CTRL or PINK1 guides. Cells in both groups were treated with 10 µM CCCP overnight. Black outline indicates significant interactors that have a log2 fold-change > 1. Proteins associated with TOM (dark blue) and TIM23 (cyan) translocases and cytosolic chaperones (yellow) are indicated. (B) Flow cytometry measurements of PINK1-YFP intensity (top) and intensity of MTS-mSc (bottom) from the same HeLaPINK1-YFP+MTS-mSc cells. N = 6 biological replicates measured from 2 independent transductions. All statistical comparisons are to the vehicle-treated CTRL guide group. **** p ≤ 0.0001. (C) Representative confocal Airyscan images shown as z-projections in HeLaPINK1-YFP cells with endogenously tagged TOMM70-Halo to mark mitochondria. Arrow indicates PINK1-YFP accumulated around lipid droplets; arrowhead indicates PINK1 co-localizing with TOMM70-Halo. Scale bar = 10 µm. (D) CLEM images demonstrating that the PINK1-YFP accumulates around lipid droplets (LD) following loss of the TOM translocase (HeLaPINK1-YFP cells). ER (yellow arrowheads) and small vesicles (red arrows) were adjacent to lipid droplet collections and accumulated PINK1-YFP. Scale bars: green = 20 µm; yellow = 5 µm; black = 1 µm; white = 500 nm. (E) Representative confocal images of TIMM23 KO pools in HeLaPINK1-YFP cells showing PINK1-YFP accumulates on mitochondria in cells lacking TIMM23 by immunostaining. Scale bar = 10 µm. (F) Line scan of dotted line in (E). (G) Quantification of (E). **** p ≤ 0.0001. N = 4 wells analyzed plated on two separate occasions from the same KO pool. Cells were analyzed 7 or 8 days after electroporation and cells without TIMM23 were scored for PINK1-YFP accumulated on TOMM40-positive mitochondria. (H) Representative immunoblots from HeLa cells with exogenous PINK1-YFP (left blot) or endogenous PINK1 (right blot) +/- 10 µM CCCP treatment for 4hrs. N ≥ 3 independent experiments. (I) Volcano plot shows PINK1-YFP (red) interactors, following CTRL vs. TIMM23 KD in HeLaPINK1-YFP cells, with protein groups color coded as in (A). N = 4 replicates/sgRNA on 1 occasion. (J) In gel fluorescence of PINK1-YFP complexes separated by CN-PAGE after mixing lysates with the indicated antibodies. Specific interaction of antibody with PINK1-YFP containing complex increases the molecular weight of the complex causing it to shift up in the gel. 10 µM CCCP treatment in indicated lanes was for 3 hrs. N = 2 independent experiments with TOMM20 gel shift, one of which also tested TOMM22 and TOMM40 antibodies.

To test which of the core components of the TOM-TIM23 supercomplex stabilize PINK1 on the OMM, we compared knockdown of TOMM22 and TOMM40 to knockdown of TIMM23 (Fig. 5B). By flow cytometry, all knockdowns caused sustained block of import, as reflected in decreased MTS-mSc levels (Fig. 5B, bottom), and all led to increased PINK1-YFP at the whole cell level (Fig. 5B, top). However, the pattern of PINK1-YFP accumulation, determined by confocal microscopy, differed dramatically following TOM vs. TIM23 perturbation. With TIMM23 KD, PINK1-YFP co-localized with the OMM marker TOMM70, consistent with our findings above (Fig. 5C, bottom). Following TOMM22 or TOMM40 KD, by contrast, most PINK1-YFP accumulated near lipid droplets, adjacent to mitochondria (Fig. 5C, top and middle, arrows, and 5D), and in the cytosol in a mixed diffuse/punctate pattern. Some weak PINK1-YFP signal co-localized with TOMM70 following TOMM22 KD but not TOMM40 KD (Fig. 5C, top, arrowhead), suggesting a more severe phenotype following depletion of TOMM40. This demonstrates that PINK1- YFP accumulates but cannot be maintained on the OMM in the absence of the TOM translocase. To verify that TOMM70 labels mitochondria in the absence of TOMM40, we performed CLEM, using a HeLa^PINK1-YFP^ cell line in which TOMM70 was additionally endogenously tagged with HaloTag (Fig. S5A). As expected, TOMM70-Halo localized to residual double membraned mitochondria in TOMM40 KD cells, in proximity to but separate from lipid droplets with PINK1-YFP accumulation.

To verify these results with an orthogonal approach we generated knockout (KO) pools for TIMM23 by electroporating Cas9 protein complexed with guide RNAs targeting TIMM23. Consistent with the findings following TIMM23 KD, PINK1-YFP accumulated on mitochondria in cells that had lost TIMM23 immunostaining (Fig. 5E – G). KO pools targeting TOMM22 and TOMM40 also showed PINK1- YFP accumulation in the same pattern as observed by CRISPRi (Fig. S5B).

Next, we evaluated PINK1 accumulation and activation following TOM and TIM23 disruption by immunoblotting. Full length PINK1-YFP accumulated following disruption of either TOM or TIM23 translocases, but only TIMM23 KD activated PINK1, as evidenced by increased pS65-Ub (Fig. 5H, left blot). This suggests that the full length PINK1-YFP accumulating in the absence of TOM is inactive.

Immunoblotting for endogenous PINK1 showed a similar pattern as exogenous PINK-YFP, except that endogenous full length PINK1 did not accumulate following TOM disruption (Fig. 5H, right blot). We hypothesize that the YFP tag may block degradation of full length PINK1, allowing PINK1-YFP but not endogenous PINK1 to accumulate following loss of TOM. In the absence of the TIM23 translocase, PINK1-YFP maintained its association with the TOM translocase by AP-MS but lost its association with TIM23 subunits (Fig. 5I and Table S5). Consistently, by CN-PAGE, PINK1-YFP was observed in a discrete complex that could be shifted to a higher molecular weight by antibodies against TOMM20, TOMM22, or TOMM40, demonstrating PINK1-YFP is bound to the TOM translocase following TIMM23 KD (Fig. 5J and Fig. S5C). Together this establishes that TOM but not TIM23 is required for PINK1 stabilization and activation on the OMM.

### PAM import motor facilitates but is not required for PINK1 import

Having established that import block at TIM23 is critical for PINK1 stabilization in the TOM complex, we next considered the role of the PAM import motor ^11^. HSPA9, an essential component of the PAM motor, was identified as a top activator in the screen, but HSPA9 has other functions in the mitochondrial matrix. These include folding newly imported proteins with the assistance of its co-chaperone DNAJA3 – which was also identified as a top activator in the screen ^39,40^.

The dual role of HSPA9 in import and protein folding has led to a proposal that misfolded proteins can activate the PINK1-Parkin pathway independently of the MMP by competing HSPA9 away from the PAM complex ^41^. To assess this proposal, we directly compared disruption of the HSPA9 - DNAJA3 complex to disruption of the HSPA9-containing PAM complex (Fig. 6A and B). KD of both PAM16 and DNAJA3 blocked import of the ATP5A precursor (Fig. 6A). However, only knockdown of DNAJA3 caused endogenous PINK1 stabilization and activation, similar to KD of TIMM23 (Fig. 6A). Using an orthogonal strategy, we compared KO pools of TIMM23 and TIMM44 on PINK1-YFP stabilization in HeLa^PINK1-YFP+MTS-mSc^ (Fig. 6C and D). In cells with blocked import of MTS-mSc, greater PINK1-YFP increase was observed with TIMM23 KO compared to TIMM44 KO (Fig. 6C and D). Similarly, PAM16 KD caused less PINK1-YFP accumulation than TIMM23 KD (Supplemental Fig. S5D and E). Thus, the PAM motor is not required for PINK1 import under endogenous conditions but may facilitate PINK1 import in the setting of higher levels of PINK1-YFP by exogenous expression.

**Figure 6.**
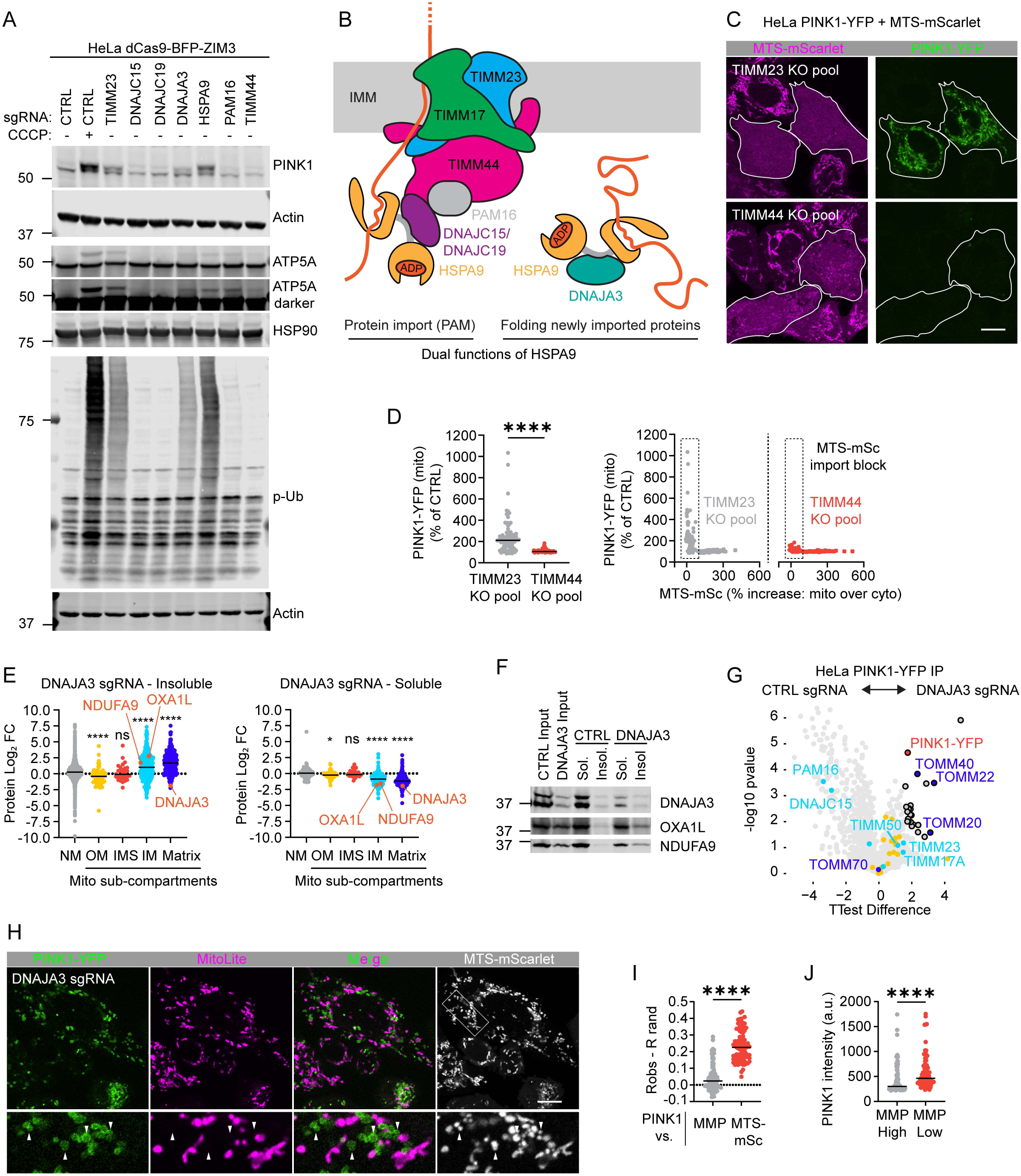
PINK1 is stabilized by protein misfolding but not disruption of the PAM import motor. (A) Representative immunoblot comparing endogenous PINK1 activation to import block of ATP5A as described for (Fig. 2J). N = 3 independent experiments. (B) Schematic showing the dual functions of HSPA9 in the PAM import motor and folding of newly imported proteins. HSPA9 performs the latter with its co-chaperone DNAJA3. (C) Representative confocal images of PINK1-YFP and MTS-mSc in TIMM23 and TIMM44 KO pools 7 days after electroporation. Chronic import block through the TIM23 translocase is identified by accumulation of MTS-mSc in the cytosol (cells with white outlines). Scale bar = 10 µm. (D) Quantification of experiment in (C). Mitochondrial PINK1-YFP intensity was measured in the subset of cells with import block (left) and correlation of PINK1-YFP with import block of MTS-mSc for all cells (right). **** p ≤ 0.0001. 89 or more cells per condition were analyzed from N = 4 wells, plated on two separate days, from the same KO pool. Cells were analyzed 7 days after electroporation. (E) LFQ proteomics depicting protein abundance from the 1% Triton X100 soluble and insoluble heavy membrane fractions, normalized to the median value of non-mitochondrial proteins in the sample. * p ≤ 0.05, **** p ≤ 0.0001. N = 4 replicates/sgRNA on 1 occasion. (F) Immunoblot of 1% Triton X100 soluble and insoluble fractions from whole cell lysates. N = 3 replicates on at least 2 occasions. (G) Volcano plot showing PINK1-YFP interactors as described in (5I). N = 4 replicates/sgRNA on 1 occasion. (H) Representative confocal image of same cell line in (Fig. 3E) transduced with a guide targeting DNAJA3. Scale bar = 10 µm. (I) Quantification of cells in (H) was performed as in (Fig. 3G). **** p ≤ 0.0001. N = 86 cells from six wells and 2 separate transductions. (J) Quantification of cells in (H) was performed as in (Fig. 3F). **** p ≤ 0.0001. N = 91 cells from six wells and 2 separate transductions.

We next examined the mechanism of PINK1 stabilization by DNAJA3 KD in greater detail.

Consistent with the function of DNAJA3 in folding newly imported proteins, DNAJA3 KD globally reduced the solubility of matrix and IMM proteins ^40^, including established substrates OXA1L and NDUFA9, by proteomics and immunoblotting in HeLa^dCas9-BFP-ZIM3^ cells (Fig. 6E and F and Table S6). This is consistent with the visible matrix aggregates by EM following DNAJA3 KD, as discussed above (Fig. 3C and Fig. S2C). Notably, PINK1-YFP stabilized by DNAJA3 KD pulled down with the TOM and TIM23 translocase subunits in a similar pattern as observed with CCCP treatment (Fig. 6G and Table S5).

We next assessed if PINK1-YFP accumulation by DNAJA3 KD is associated with MMP loss.

Consistent with single cell analysis by flow cytometry, PINK1-YFP was elevated selectively on mitochondria with low MMP by confocal microscopy in HeLa^PINK1-YFP^ ^+ MTS-mSc^ cells (Fig. 6 H – J). This followed the same pattern as NDUFAB1 and PMPCB and was distinct from the pattern resulting from direct import block with TIMM23 KD (Fig. 4H – J). Together these findings suggest that misfolded proteins from DNAJA3 KD activates the PINK1-Parkin pathway by disrupting the MMP. The PAM motor, by contrast, is dispensable for import of endogenous PINK1.

### TOMM20 and TOMM5 are required for PINK1 stabilization on the OMM, while TOMM70 is dispensable

Our findings so far suggest that TOM but not the TIM23 translocase is required for PINK1 stabilization on the OMM, and that the MMP is the primary driving force for PINK1 import through the TIM23 translocase. We next explored what specific subunits of the TOM translocase are required to hold PINK1 on the OMM, in addition to the core subunits TOMM40 and TOMM22 (Fig. 5C). The TOM translocase has two receptors for mitochondrial precursors in addition to TOMM22: TOMM20 and TOMM70 ^42^.

Consistent with recent reports ^5,43^, we found TOMM20 KD blocks PINK1 stabilization following OXPHOS inhibition (Fig. 7A, left, and 7B, left blot, lane 4 vs. 2). Notably, however, TOMM70, which is required for PINK1 import in *in vitro* import assays and in yeast ^43,44^, was not required for PINK1 import in intact mammalian cells (Fig. 7A, left, and 7B, left blot, lane 6 vs. 2). This is consistent with the absence of binding between PINK1-YFP and TOMM70 on de-energized mitochondria (Fig. 5A).

**Figure 7.**
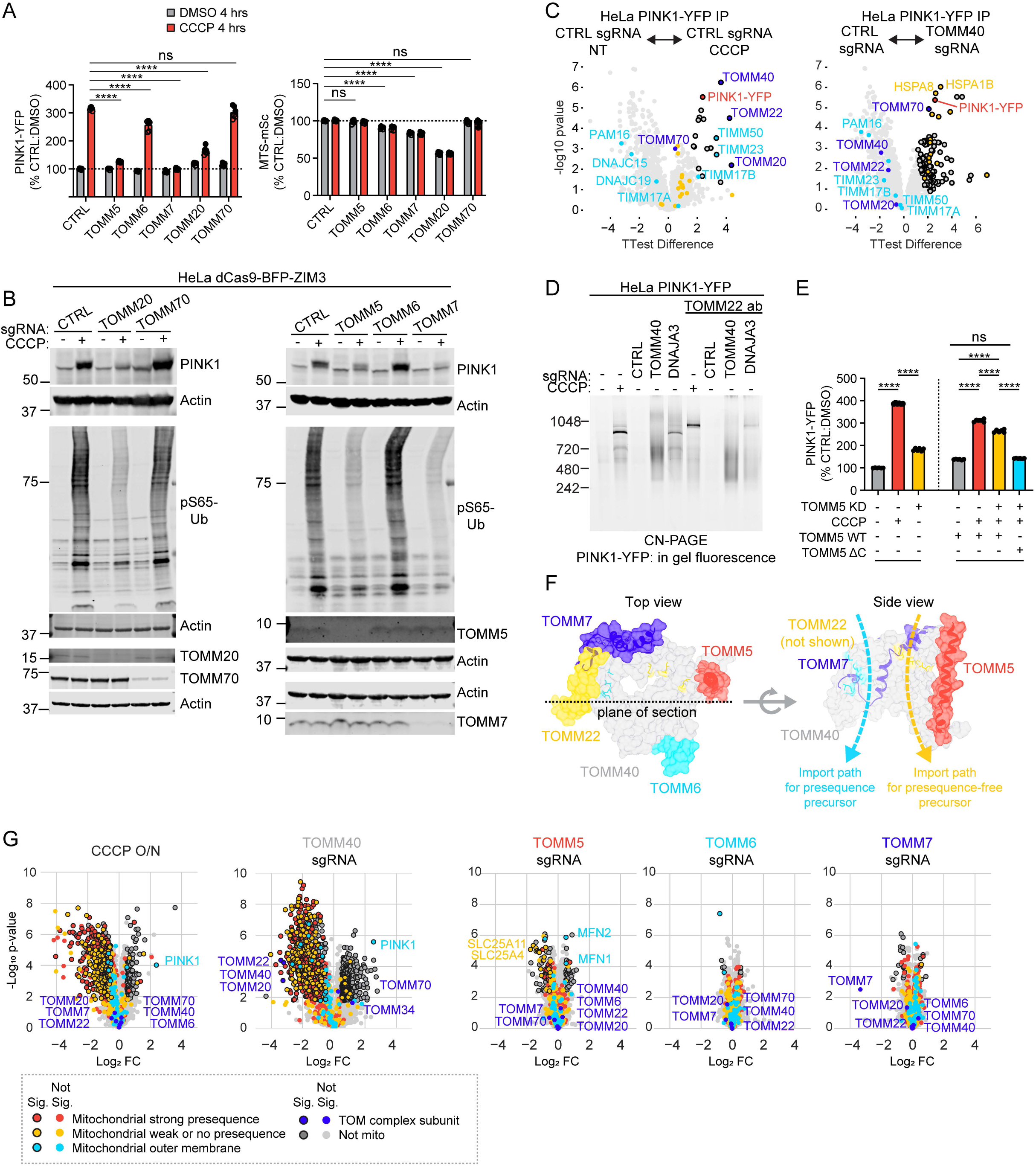
TOMM20 and TOMM5 are required for PINK1 stabilization on the OMM; TOMM70 is not. (A) Flow cytometry measurements performed as in (Fig. 5B). **** p ≤ 0.0001. Error bars mean +/- SD. N = 6 independent experiments from 2 separate transductions. (B) Representative immunoblots of HeLadCas9-BFP-ZIM3 cells +/- 10 µM CCCP for 4 hrs. N ≥ 3 independent experiments. (C) Volcano plots of PINK1-YFP interactors measured by AP-MS as in (5I). CCCP treatment, where indicated, was overnight. Other samples were untreated. N = 4 replicates/sgRNA on 1 occasion. The untreated control guide group was the same as used in (Fig. 6G). (D) CN-PAGE separated PINK1-YFP complexes visualized by in gel fluorescence as in (Fig. 5J). N = 2 independent experiments with TOMM40 KD, one of which was with antibody gel shift. (E) Flow cytometry in HeLaPINK-YFP cells with doxycycline inducible expression of wildtype or C-terminal truncated TOMM5. N = 6 replicates on at least 2 occasions. **** p ≤ 0.0001 (F) Crystal structure yeast TOM complex (PDB: 6JNF 45) demonstrating location of TOM5 and TOM7 subunits. (G) LFQ proteomics of HeLaPINK-YFP whole lysates following the indicated knockdown/treatment. N = 4 replicates/sgRNA on 1 occasion.

Surprisingly, however, we found that PINK1-YFP did bind TOMM70 when the TOM translocase was depleted by TOMM40 KD (Fig. 7C and Table S5). Under these conditions, PINK1-YFP additionally pulled down cytosolic HSP70 chaperones and co-chaperones, which are well-established binding partners of TOMM70 ^42^ (Fig. 7C and Table S5). Consistently, by CN-PAGE, most of the PINK1-YFP in TOMM40 KD cells was found in a smear between 480 and 720 kDa that did not shift following incubation with an antibody against TOMM22, in contrast to PINK1-YFP stabilized by CCCP treatment or DNAJA3 KD (Fig. 7D and Fig. S5F). Together, these data suggest that TOMM70 may bind excess unfolded PINK1 that cannot be incorporated into the TOM translocase or degraded in the cytosol. At the same time, they demonstrate that TOMM70 is not required for endogenous PINK1 import or stabilization in the TOM translocase.

Finally, we considered the remaining three small subunits of the TOM complex, TOMM5, TOMM6, and TOMM7 ^42^ (Fig. 7A, B, E, and F). TOMM7 has previously been shown to be critical for PINK1 stabilization in the TOMM40 complex ^18^, however, the mechanism has not been clear. We confirmed that TOMM7 is required for PINK1 stabilization on the OMM, but, surprisingly, found that TOMM5 is also needed (Fig. 7A, left, and 7B, right blot, lane 4 vs. 2 and lane 8 vs. 2). Stabilization of PINK1-YFP could be restored by exogenous expression of full length TOMM5 but not C-terminally truncated TOMM5 (Fig. 7E). Notably, TOMM5 and TOMM7 were not required for stability of the core subunits TOMM40 or TOMM22 or the receptor TOMM20, by quantitative proteomics (Fig. 7G, blue data points and Table S7). Consistently, neither TOMM5 nor TOMM7 was required for import of presequence-containing proteins through the TOM translocase (Fig. 7G, red data points, and Table S7). This was in sharp contrast to TOMM40 KD or import block with CCCP overnight, which strongly depleted presequence-containing mitochondrial proteins (Fig. 7G, red data points, and Table S7). In the previously published TOM structures, TOMM5 and TOMM7 are not in direct contact, but are predicted to stabilize the interior of the translocase pore, with TOMM5 helping to shape the import path for hydrophobic precursors lacking an import presequence (Fig. 7F) ^45^. Consistently, the abundance of presequence-less mitochondrial proteins was reduced following KD of TOMM5 (but not TOMM6 or TOMM7) (Fig. 7G), demonstrating functional conservation between mammalian TOMM5 and its yeast ortholog ^45^. Together these findings suggest that TOMM5 and TOMM7 may stabilize a binding site for PINK1 in the translocase overlapping with the import path for proteins lacking a precursor sequence. This binding site is likely separate from the previously identified binding site between PINK1 and TOMM20 ^5,43^, as TOMM20 was not destabilized by loss of the small TOM subunits.

## Discussion

Here, we used a novel single-cell reporter for the PINK1-Parkin pathway, MFN2-Halo, to identify facilitators and activators of the PINK1-Parkin pathway in genome-wide screens. Notably, functionally diverse mitochondrial perturbations, including those causing severe mitochondrial protein misfolding, converged on loss of MMP as the primary mechanism of PINK1-Parkin activation. Consistently, we established that the MMP is the main driving force for PINK1 import through the TIM23 translocase, with mitochondrial ATP and the PAM motor playing supporting roles.

Surprisingly, TIMM23 KD was among the novel PINK1-Parkin activators identified. In contrast to the other activators, TIMM23 KD stabilized PINK1 on energized mitochondria in complex with the TOM translocase, demonstrating that block of transport through the TIM23 translocase is the key step for PINK1 stabilization and activation. We further identified components of the TOM translocase required for PINK1 stabilization on the OMM, including a novel role for TOMM5, which may stabilize a PINK1 binding site in the TOM translocase distinct from the established TOMM20 binding site. Finally, we identified that the PINK1-Parkin response is additionally shaped by the overall energetic state of the cell due to the high-energy requirement for new PINK1 synthesis. Together these findings suggest a refined model for damage-sensing in the PINK1-Parkin mitophagy pathway: the PINK1-Parkin pathway is triggered when the force of the MMP across the TIM23 translocase can no longer overcome the binding free energy that holds PINK1 in the TOM complex.

This model helps clarify several open questions in the field. Mitochondrial uncoupling with CCCP was shown to be sufficient to activate the PINK1-Parkin pathway over a decade ago ^22,46^, leading us and others to propose that activation of the PINK1-Parkin pathway is due to import block of PINK1 along the precursor pathway ^47^. This model, however, was seemingly at odds with recent studies, which found TIMM23 knockdown by transient siRNA mediated KD (for 2 – 3 days) or small molecule inhibitors does not phenocopy PINK1 stabilization by CCCP ^4,48,49^. Using methods that allowed for more complete TIMM23 knockdown or knockout in single cells (assayed at 7 - 8 days after transduction or electroporation), we found that TIMM23 depletion is sufficient to stabilize and activate PINK1. This clarifies that import block at TIM23 translocase is a key step in PINK1 stabilization on the OMM and reinforces the import block model of PINK1-Parkin activation.

Our results additionally help clarify which components of the recently identified PINK1-TOM- TIM23 supercomplex are required for PINK1 stabilization on the OMM ^4,5^. While we confirm that PINK1 binds both TOM and TIM23 translocases, we found only the TOM translocase is required for PINK1 stabilization on the OMM. Stable contacts with the TIM23 translocase, while dispensable for PINK1 stabilization on the OMM, may enable rapid import of PINK1 to the IMM for cleavage, if the MMP is restored ^29^, and may additionally protect import arrested PINK1 from intermembrane space proteases such as OMA1, as recently proposed ^4^. Among the TOM subunits required for PINK1 stabilization, we confirmed that previously identified subunits TOMM40, TOMM22, TOMM20, and TOMM7 are essential ^5,18,43^, and additionally identified an essential role for TOMM5. By contrast, we found that TOMM70 was not required for endogenous PINK1 import but bound excess PINK1-YFP when the TOM translocase was depleted. This likely explains why prior studies found that TOMM70 is required for exogenous PINK1 import in *in vitro* assays and following exogenous PINK1 expression in yeast ^43,44^. In these assays, exogenous PINK1 may exceed the capacity of the precursor pathway and reversibly bind cytosolic HSP70 chaperones and TOMM70.

Another open question concerns the driving force for PINK1 import along the precursor path: is only the MMP required or is the PAM import motor also needed? This question has been made more relevant by a recent proposal that the PAM motor may “sense” protein misfolding in the mitochondrial matrix to activate the PINK1-Parkin pathway independently of the MMP ^41^. This mechanism assumes that that the PAM motor is essential for PINK1 import, but this assumption had not been directly tested ^41^. Here, we identified that the MMP is the primary driving force for PINK1 import across the TIM23 translocase, while the PAM motor is not required for endogenous PINK1 import. The greater dependence of PINK1 on MMP than PAM for import is supported by prior work in yeast: import of single-pass proteins to the IMM, which involves translocation of only a short N-terminal stretch of the positively charged amino acids across the IMM, is less dependent on the PAM motor than precursor import to the mitochondrial matrix, involving a substantially longer and more varied stretch of amino acids ^11^. Consistently, we found that import of the matrix targeted proteins ATP5A1 and MTS-mSc were more sensitive to disruption of the PAM import motor.

Importantly, the proposed model also points to how the PINK1-Parkin pathway may be pharmacologically tuned to promote clearance of damaged mitochondria in disorders caused by mitochondrial damage. Small molecules that upregulate glycolysis, such as the PGK1 activator terazosin ^50–52^, may help support PINK1-Parkin surveillance of mitochondrial damage, by providing ATP needed for new PINK1 synthesis and PINK1-Parkin activation. This may be particularly critical in sporadic Parkinson’s disease, where energy deficiency is likely present in the affected dopamine neurons ^52^. Additionally, the MMP threshold of PINK1-Parkin activation may be lowered by decreasing mitochondrial ATP levels. This may have an effect that is similar to MTK458, a PINK1 activator under development by Mitokinin/AbbVie, which was shown to lower the CCCP dose required for PINK1-Parkin activation ^53^, similar to the effect of oligomycin observed here. Conversely, small molecules that stabilize the binding site for PINK1 in the TOM complex, buttressed by TOMM5 and TOMM7, may allow PINK1 stabilization at a higher MMP. Either strategy – strengthening PINK1 binding to the TOM translocase or lowering the driving force through the TIM23 translocase – would have a similar effect: increasing the sensitivity of the PINK1-Parkin mitophagy pathway for damaged mitochondria. Finally, we anticipate the MFN2-Halo reporter will help aid drug discovery, as a quantitative, single-cell reporter that is orthogonal to the widely used mitophagy-based reporters.

## Resource availability

### Lead contact

Further information and requests for reagents will be fulfilled by Lead Contact Derek Narendra.

### Materials availability

All constructs or cell lines generated in this study are available from the lead contact upon request and completion of a Material Transfer Agreement.

### Data code and availability

- Raw sequencing data from the screens will be submitted to the NCBI’s Gene Expression Omnibus (GEO) Database (accession numbers: TBD).
- Original code will be made available on (https://github.com/NarendraLab/Code-used-in-Thayer-JP-et-al.-2025).
- Any additional information required to reanalyze the data reported in this paper is available from the lead contact upon request.

## Acknowledgements

We thank Dr. Dragan Maric and the NINDS Flow Cytometry Facility for technical assistance with FACS, Dr. Carolyn Smith and the NINDS Light Microscopy Facility for technical assistance with confocal microscopy, and Dr. Yuesheng Li and the NHLBI DNA Sequencing and Genomics Core for technical assistance with next-generation sequencing. This work utilized the computational resources of the NIH HPC Biowulf cluster (https://hpc.nih.gov). We thank Dr. Kei Okatsu (Kyoto University) for technical advice on in gel imaging of PINK1-YFP by CN-PAGE. We thank Dr. Richard Youle for his critical reading of the manuscript and insightful comments. This work was supported by the Intramural Research Program of the NINDS, National Institutes of Health (grant: 1ZIANS003169).

## Author contributions

J.A.T. and D.P.N. led the project. J.A.T., D.P.N., J.D.P, X.H., and Y.L. designed and performed experiments.

J.H., D.M.R., S.S. and M.E.W. supported the project by providing supervision, suggestions, and reagents.

J.A.T. and D.P.N. wrote an initial draft of the manuscript. All authors edited the manuscript.

## Supplemental information

Figures S1 – S5.

Tables S1 – S10. Excel files containing additional data too large to fit in a PDF.

## STAR Methods

### Cell Culture

All HeLa and HEK293 cells were grown in high glucose DMEM with pyruvate (Gibco, 11995-065) supplemented with 10% fetal bovine serum (R&D Systems, S11550) and 1% Penn/Strep (Gibco, 15140-122) and maintained in an incubator at 37⁰C and 5% CO2, except where indicated. For experiments to test alternative carbon sources (galactose, pyruvate, low glucose, and no glucose) on PINK1 stabilization, DMEM without glucose and pyruvate but with L-glutamine (Gibco, 11966025) was supplemented with the indicated carbon source. The following lines were used throughout the manuscript to generate new cell lines described below. HeLa dCas9-BFP-KRAB ^54^ (gift of the Dr. Richard Youle Lab), HeLa PINK1 KO PINK1-eYFP Clone #21 ^48^ (from the Dr. Shiori Sekine Lab), WT HeLa (ATCC), WT HEK293 (ATCC).

#### Generation of cell lines

dCas9-BFP-ZIM3 was introduced to cell lines via the piggybac system - iC30 PB-zim3-mycNLS-BFP-hygro- ucoe (iNDI iPSC Neurodegenerative Disease Initiative) and K13-EF1a-transposase. Cells were transfected with the transposase and the dCas9-BFP-ZIM3 plasmid with FuGENE Transfection Reagent (HEK293 cells - FuGENE 6 Promega, E2691 and HeLa cells – FuGENE HD Promega, E2311) following manufacturer’s instructions. Briefly, for HeLa cells a ratio of 3:1 FuGENE HD was used (6µL:2µg) and for DNA 1:2 ratio transposase:dCas9. For HEK293 cells a ratio of 4:1 FuGENE 6 was used (8µL:2µg) and for DNA 1:2 ratio transposase:dCas9. Once cells were expressing BFP the cells were selected with hygromycin 200µg/mL (MP Bio, 194170) for 72hrs. Finally, cells were sorted via FACS for a clonal population which was expanded and used for experiments. HeLa dCas9 BFP ZIM3 was added to WT HeLa cells, HeLa PINK1 KO PINK1 EYFP (Clone #21 From Dr. Shiori Sekine Lab), and WT HEK293 cells.

To generate stable lines, cells were transduced with the indicated virus and 8µg/mL polybrene. Following transduction cells were sorted via FACS to obtain a homogeneous population. Retroviral or lentiviral vectors were used encoding mCherry-Parkin (gift from Dr. Richard Youle Lab), YFP-Parkin (gift from Dr. Richard Youle Lab), MTS(Cox8a)-mScarlet, and/or mt-Keima (Gift from Dr. Richard Youle Lab).

Endogenously tagged cell lines were made by nucleofection according to the manufacturer’s instructions. MFN2 was endogenously tagged in HeLa dCas9-BFP-KRAB and HEK293 dCas9-BFP-ZIM3 cell lines. TOMM70 was endogenously tagged in the HeLa PINK1 KO PINK1-eYFP dCas9-BFP-ZIM3 cells.

Briefly, guides targeting the protein of interest and donor plasmids (that included the Halo tag sequence as well as 1kB homology arms) were added together with HiFi Cas9 Nuclease V3 (IDT, 1081061) along with the nucleofection reagent. For HeLa cells the SE Cell Line 4D-Nucleofector X Kit L (Lonza, V4XC- 1012) and for HEK293 cells the SF Cell Line 4D-Nucleofector X Kit L (Lonza, V4XC-2024). The mix was added to cells and nucleofected via the Lonza 4D-Nucleofector Core Unit (Lonza, AAF-1003B) and 4D- Nucleofector X Unit (Lonza, AAF-1003X) nucleofector. Program for HeLa cells CN-114 and for HEK293 cells CM-130. The cells were then plated in media containing HDR enhancer (IDT, 1081073 or IDT, 10007921). The next day media was replaced with media not containing HDR enhancer. Cells were sorted via FACS and expanded to use for further experiments. (See material list for guide sequences).

Nucleofection was also used to generate cells for the KO pool experiments. Briefly, the same protocol was completed as detailed above – Cas9 protein and sgRNA were combined to form a ribonucleoprotein complex in vitro which was electroporated into the cells – but there was no donor plasmid added or HDR enhancer. Cells were analyzed at least 7 days after electroporation. (See material list for guide sequences).

#### Drug treatments and reagents

HeLa or HEK293 cells were treated with the following drugs/concentrations where indicated oligomycin 10 µg/mL (Sigma-Aldrich, Inc 75351-5MG), antimycin A 4 µg/mL or 8 µg/mL (Sigma-Aldrich, Inc, A8674), 2-deoxy-D-glucose (2-DG) 10 mM (Sigma-Aldrich, D8375-5G), carbonyl cyanide 3- chlorophenylhydrazone (CCCP) 10 µM (Sigma, C2759-100MG), MG132 50 µM (Sigma, SML1135), cycloheximide 50 µg/mL (Sigma-Aldrich, Inc, C7698-5G), heptelidic acid 10 µM (Cayman chem, 14079), rotenone 1 µM (Calbiochem, 557368-1GM), KL11743 10 µM (Millipore Sigma, SML3458-25MG).

Day 10 i^3^Neurons were treated with CCCP 20 µM where indicated.

When appropriate Janelia Fluor 646 HaloTag Ligand (Promega Corporation, GA1121) was added to the cells at 75 nM (based off titration experiments) for 25 minutes before analyzing.

#### Lentivirus production

Lentivirus was produced in Lenti-X cells. FuGENE 6 Transfection Reagent (Promega, E2691) following manufacturer’s instructions was used to generate individual viruses. Briefly, Lenti-X cells were transfected using a ratio of 3:1 FuGENE 6:DNA (6µL:2µg) the day after cell seeding. For a 6 well plate the cells were transfected with packaging plasmids psPAX2 (750ng) and pMD2.G (250ng) as well as the appropriate lentiviral plasmid (1µg). Media was removed the following day and replace with 1mL. 72-96 hrs post transfection media was collected and centrifuged at 3400g, 10 minutes, 4⁰C. Being careful to not disturb the pellet, supernatant was moved to a new tube, aliquoted, and froze at -80⁰C.

To make virus of the dual-sgRNA library from the Weissman Lab (addgene, 187246) Lipofectamine 3000 (Invitrogen, L3000001) was used following manufacturer’s instructions ^55^. Briefly, cells were plated on poly-L-ornithine (PLO) (Sigma, P3655) coated 15cm plates. The next day cells were transfected with packaging plasmids (psPAX2 (13.3µg), pMD2.G (4.5µg), and pAdvantage (1.8µg)) as well as with the lentiviral dual-sgRNA plasmid (19.5µg), 60uL of Lipofectamine 3000 reagent and 80uL P3000 enhancer reagent. Full media change was completed the next day and media was collected 96 hrs post transfection. Virus was concentrated with Lenti-X Concentrator (Takara Bio, 631231) following manufacturer’s instructions. Briefly, media was collected and centrifuged at 3400g, 10 minutes, 4⁰C. Being careful to not disturb the pellet, supernatant was moved to a new tube and Lenti-X Concentrator was added at a 1:3 ratio. The media/concentrator mix was placed at 4⁰C for 24 - 48 hrs. Next, the solution was pelleted at 1500g, 45 minutes, 4⁰C. The supernatant was discarded and the pellet was resuspended in PBS at 1/10 of the original volume of media collected off the plate. Virus was aliquoted and frozen at -80⁰C.

#### siRNA Transfections

The indicated cell lines were transiently transfected utilizing Lipofectamine RNAiMax Reagent (Life Technologies, 13778) per manufacturer’s instructions with 20 nM final concentration of siRNA. siRNA used –ON-TARGETplus SMARTPool Human PINK1 (Dharmacon, L-004030-00-0005) and siGENOME Non-Targeting siRNA #1 (Dharmacon, D-001210-01-05).

#### Flow cytometry

All flow cytometry data was acquired on a Cytek Amnis CellStream Flow Cytometer Four-Laser System with 405 nm, 488 nm, 561 nm, and 642 nm and AutoSampler (Cytek, CS-100496). Cells were assessed live, except for one experiment to demonstrate retention of MFN2-Halo signal in fixed samples. For that experiment, HeLa^MFN2-Halo^ ^+ mCh-Parkin^ cells were incubated in fixation buffer (BD Biosciences, 554655) for 10 minutes followed by PBS washes. The cells were immediately analyzed by flow cytometry or stored at 4⁰C in PBS for one day and then analyzed.

For all flow cytometry experiments using dual guides, a population gated for single cells was further gated for cells expressing guide (TagBFP2+) and cells not expressing guide (TagBFP2-).

For analysis of MFN2-Halo degradation among the PINK1-Parkin activators and facilitators, MFN2-Halo in the mCh-Parkin positive (Parkin+) population was compared to the mCh-Parkin negative (Parkin-) population within each sample to determine Parkin specific changes to MFN2-Halo intensity following each knockdown. Specifically, the raw intensity of each population, Parkin+ and Parkin-, was first normalized by dividing by the average raw intensity of CTRL guide samples, giving MFN2-Halo normalized intensity values (MFN2normint)(except in figure 1E where cells were not gated on mCh-Parkin). The percent MFN2-Halo degradation was then calculated as follows:

MFN2 degradation (%) = 100 * ((MFN2normint^Parkin+^ - MFN2normint^Parkin-^) / MFN2normint^Parkin+^.

For analysis of PINK1-YFP, the raw YFP intensity for guide positive cells was normalized by dividing by the average of the CTRL replicates in the sample. Similarly, for analysis of MTS-mSc, the raw mScarlet intensity for guide positive cells was normalized by dividing by the average of the CTRL replicates in the sample. For experiments using TMRE to measure MMP, cells were incubated with TMRE 20 nM for 15 – 20 minutes prior to trypsinization for cell sorting. The raw TMRE intensity was normalized to the untreated condition.

Mitophagy was measured with the mt-Keima reporter by plotting the ex. 488 - em. 611/31 channel (representing neutral mKeima) vs. ex. 561 – em. 611/31 (representing acidic mKeima). To compensate for bleed-through from the YFP-Parkin channel, 10% compensation was applied for the ex. 488 – em. 528/46 channel into the ex. 488 - em. 611/31 channel. A triangular gate was drawn to capture events that shifted with mitophagy into a high acidic/neutral population. The percent of cells in the “mitophagy” gate was reported.

For experiments measuring both PINK1-YFP and MMP, the cells were incubated for 15 – 20 minutes with MitoLite NIR following the manufacturer’s instructions prior to sorting. We found the raw intensity of the MitoLite NIR staining from well-to-well was sensitive to cell number. To control for well-to-well variability, we normalized the MitoLite NIR signal as follows: (1) the raw values for the BFP+ population was divided by the raw value for the BFP- population to obtain within-sample normalized values, and then (2) the normalized values for each guide were divided by the average of the normalized values for the CTRL guide samples in the same experiment.

For 2D kernel density plots comparing PINK1-YFP and MMP (measured with MitoLite NIR), analysis was performed using a custom Python script and the Pandas and NumPy libraries. To control for variability in MitoLite NIR staining from sample to sample, the average MitoLite NIR intensity for events in the PINK1- YFP low / BFP low population for each sample was calculated. These represented non-transduced cells within each sample (“MMP guide negative”). The ratio of the “MMP guide negative” MitoLite NIR intensity of each sample was divided by that of the CTRL guide sample to give a “normalization factor” for each sample. The MitoLite NIR intensity for events in each sample were divided by this normalized factor. PINK1-YFP intensity values were not modified. The Seaborn library was used to plot a 2D kernel density plot for each sample. Events in each of the four quadrants were calculated using the Python script using the same cutoff values for each sample. Stacked bar graphs of these data were generated in Graphpad Prism 10.

#### SUNset Assay

Cells were treated with 10 µM Puromycin (Invivogen, ant-pr-1) for 10 minutes before immediately lysing the cells. Lysates were run via western blot and probed with puromycin antibody (see immunoblotting section).

#### iPSC Culture and Neuronal Differentiation

i11W-mNC iPSC were used for all neuronal differentiations (these cells are the WTC11 iPSCs expressing doxycycline-inducible mouse neurogenin-2 (NGN2) and CAG-dCas9-BFP-KRAB). Cells were maintained and differentiated to i^3^Neurons according to published protocols ^56^. Briefly, iPSCs were plated on Matrigel (CORNING, 354277) coated plates and cultured in Essential 8 Medium + Supplement (Gibco, A1517001). Following a thaw or a passage 10 µM Y-27632 ROCK inhibitor (Tocris Bioscience, 1254) was added to the media to promote survival. Passaging was done with 0.5mM EDTA (Invitrogen, 15575-038) or when single cells were needed with accutase (Gibco, A1110501). For differentiation, cells were dissociated via accutase and plated in induction medium consisting of KnockOut DMEM/F12 (Gibco, 12660012), N2 Supplement 100X (Gibco, 17502048), non-essential amino acids 100X (Gibco, 11140050), L-glutamine 100X (Gibco, 25030081), Y-27632 ROCK inhibitor (Tocris Bioscience, 1254), doxycycline (Sigma, B9285). Complete media changes were done for the next 2 days, on day 3 cells were plated to poly-L-ornithine (PLO) (Sigma, P3655) coated plates. Cells were plated in cortical neuron culture media consisting of BrainPhys neuronal medium (STEMCELL Technologies, 05790), B27 supplement (Life Technologies|AB / Invitrogen, 17504044), BDNF (PeproTech, Inc., 450-02-50UG), NT-3 (PeproTech, Inc., 450-03-50UG), Laminin (Life Technologies|AB / Invitrogen, 23017015), and doxycycline. Half media changes were performed on the cells every other day until collection day.

#### Genome-wide CRISPRi Screen

The genome-wide CRISPRi screens were all performed in duplicate. Cells were transduced with the dual-sgRNA library from the Weissman Lab (Dual sgRNA CRISPRi Library 1-2 was a gift from Jonathan Weissman Addgene #187246) at an MOI of 0.3 based off the virus titer ^55^. Puromycin selection was added to the cells (2.5µg/mL) 48 hrs post transduction for 2 days followed by 3 days of no puromycin media. For HeLa cells, AO was added for 4 hrs 40 minutes, and for HEK293 cells CCCP treatment was added overnight. Each condition was performed in duplicate. The day of the screen sort, Halo Ligand 646 (Promega Corporation, GA1121) was added to the cells for 25 minutes pre-collection at 75 nM. Cells were collected and resuspended in cell sorting media (145 mM NaCl, 5 mM KCl, 1.8 mM CaCl_2_, 0.8 mM MgCl_2_, 10 mM Hepes, 10 mM glucose, 0.1% BSA). Finally, the cells were sorted on a MoFlo Astrios flow cytometer (Beckman Coulter, B52102) using Summit software. Two populations were collected representing the top 30% and bottom 30% MFN2-Halo expressing cells (6 million cells were collected per bin). After sorting, cells were spun down at 400g for 5 minutes and pellets were frozen at -80⁰C. Genomic DNA was extracted using the NucleoSpin® Blood Genomic DNA from blood kit (Macherey- Nagel™, 740954.100) according to manufacturer’s instructions. Briefly, cell pellets were thawed at room temperature, washed, and resuspended in 2mL PBS with 150µL Proteinase K. 2mL of BQ1 was added to each sample and vortexed vigorously, followed by a 15 minutes incubation at 56⁰C. Samples were cooled at room temperate for 1 hour. 2mL of 100% ethanol was added to each sample and immediately inverted to mix. Samples were loaded onto the columns provided in the kit in 2 stages (3mL for the first spin and then the rest of the sample for the second spin). Samples were spun at 4500g for 3 minutes and 5 minutes respectively. Membranes were washed twice with 2mL BQ2 at 4500g for 2 minutes and 10 minutes, respectively. Finally, DNA was eluted with pre-warmed Buffer BE. 100µL of Buffer BE was added to the membrane, incubated for 2 minutes, and spun at 4500g for 2 minutes. This was repeated with another 100µL of Buffer BE. PCR was done to amplify the region of interest/add barcodes/Illumina adaptors, as described ^55^. PCR reactions were made up of 10µg of genomic DNA, 100µM of each primer (Table S8), and NEBNext Ultra II Q5 Master Mix (New England Biolabs, M0544). The PCR conditions were as follows, 98⁰C for 30 seconds, 19 cycles of 98⁰C for 10 seconds and 66⁰C for 75 seconds, then 72⁰C for 5 minutes, and 4⁰C hold. Finally, the samples were purified with 0.5X-0.65X SPRI beads (Beckman Coulter, B23318) cleanup. 150 µL of beads was added to 300µL of PCR reaction, mixed and incubated at room temperature for 10 minutes. Beads were separated from samples on a magnetic stand for 5 minutes and the supernatant was moved to a new tube. Next 45µL beads was added to the supernatant, mixed, and incubated for 10 minutes at room temperature. Tubes were placed on a magnetic stand for 5 minutes and the supernatant was disposed of. Beads were washed for 2 minutes with 1mL fresh 80% ethanol 2 times. Finally, cells were air dried for 3 minutes and eluted in 30uL of elution buffer ^55^. PCR size and quality was verified by TapeStation. Libraries were pooled and sequenced on a NovaSeq 6000 V1.5 with a 15% PhiX spike, SP 100 cycle kit, targeted seq, PE-50-10-10-50, with Illumina and custom primers. Primers: Read 1- gtgtgttttgagactataagtatcccttggagaaccaccttgttgG Read 2- tgctatgctgtttccagcttagctcttaaac Index i5 Primer- acagttagggtgagtttccttttgtgct. Samples were demultiplexed using i5 index. FASTQ files were analyzed using MAGeCK-Vispr robust ranked algorithm pipeline to compare high 30% vs. low 30% MFN2-Halo fluorescence ^57^. The log 2-fold change values and uncorrected two-sided p-values for individual guides in the sgrna summary output file was used for further analysis, using a custom Python script. Guides with 0 read counts in one of the replicates were filtered out. Bonferroni corrected p-values were corrected by dividing the uncorrected p-values by the total number of guides. Gene scores were calculated as the product of the -log 10 corrected p-values and the log2 fold change. Genes were annotated as mitochondrial based on their inclusion in MitoCarta3.0 ^58^. The following gene names were modified prior to annotation with MitoCarta3.0: “TIMM23B” was changed to “TIMM23”; “HSPE1-MOB4” was changed to “HSPE1”; “PARK2” was changed to “PRKN”. “Parkin activators” were significant and had a log2 fold change of < -1 in not treated HeLa cells with mCherry-Parkin. Additionally, the difference in log2 fold change between the HeLa cells with mCherry-Parkin and without Parkin was < -1.25. “Parkin facilitators” were significant and had a log2 fold change > 1 in HeLa cells with mCh-Parkin that were treated with oligomycin and antimycin. Additionally, the log2 fold change was at least greater than 1 compared to log2 fold change from the Parkin mCh-Parkin untreated samples and the log2 fold change from HeLa no Parkin cells treated with oligomycin and antimycin. “MFN2 downregulators” were significant and had a log2 fold change less than -1 in both the HEK293 untreated and HeLa no Parkin untreated comparisons. Additionally, they were not “Parkin activators”. “MFN2 upregulators” were significant and had a log2 fold change greater than 1 in both the HEK293 untreated and HeLa no Parkin untreated comparisons. Additionally, they were not “Parkin facilitators”.

#### Guide Transductions

pJR103 plasmid (Addgene plasmid # 187242 ; http://n2t.net/addgene:187242 ; RRID:Addgene_187242) was digested with XbaI and XhoI and ligated with “empty vector fragment” via NEBulider (according to manufacturer’s instructions). Briefly, 50 ng vector was ligated with “empty vector fragment” at a 2:1 insert:vector ratio. This new plasmid, lenti_DualsgRNAEmptyVector_Puro_T2A_BFP2, was then digested with BsmBI and NEBuilder reactions (same amount/ratio as above) were done with the indicated sgRNA gene block fragments (Table S9). Ligations were transformed into ONE SHOT STBL3 COMP E COLI (Life Technologies, C737303) according to manufacturer’s instructions. Individual colonies were picked and grown up in LB broth, mini prepped with the QIAprep Spin Miniprep Kit (Qiagen, 27106), and sequenced (sequencing primer - ggcttaatgtgcgataaaagacaga). Lentivirus was generated for each sgRNA (see above Lentivirus production section). On the day of transduction, sgRNA was added to cells with 8µg/mL polybrene (Millipore Sigma, TR-1003-G). 48 hours post transduction media was changed to include puromycin 2.5µg/mL for 2 days, cells were then allowed to recover in non-puromycin media and were treated and collected on day 7 post transduction (unless otherwise indicated).

#### Immunoblotting

Sample prep – Cells were collected in PBS (Gibco, 10010-031) and lysed in RIPA buffer (CST, 9806) containing PI/PS (CST, 5872), sonicated (QSonica Q800R3, Q800R3-110) and a BCA assay (Thermo Scientific, 23225) was performed. Lysates were diluted in 4X buffer (BIO-RAD, 1610747) with 2- Mercaptoethanol (BME) (BIO-RAD, 1610710) and ran on Criterion TGX Precast Gels 7.5% (Bio-Rad, 5671024) or 4-15% (Bio-Rad, 5671084). Gels were transferred via the Bio-Rad TransBlot Turbo Transfer System (BIO-RAD, 1704150) onto nitrocellulose membranes (BIO-RAD, 1704272). Blocked in 5% nonfat milk (dot scientific inc., DSM17200) and incubated with primary antibody overnight at 4⁰C. The following day appropriate secondary antibodies were added before imaging the blots on a LI-COR Odyssey CLx Imager. Any western blot quantification was done using Image Studio Lite Version 5.2.

To probe for endogenous PINK1, which is present in cells at a low protein copy number, we followed a different procedure. Cells were directly lysed in 1X sample buffer containing BME and PI/PS. Samples were run on a 7.5% gel, transferred using the Biorad system and blocked in Licor PBS blocking buffer (LI-COR, 927-70001). Membranes were incubated in PINK1 primary antibody for 2 days at 4⁰C before secondary was added and membrane was imaged.

Primary antibodies – PINK1 (CST, 6946S), β-Actin (Sigma, A2228), Parkin (CST, 2132S), HSP90 (Proteintech, 13171-1-AP), MFN2 (Abcam, ab56889), β-Tubulin (Sigma, T8328), pUb S65 (CST, 62802S), Puromycin (Sigma, MABE343), ATP5A (Abcam, ab14748), ENO1 (Proteintech, 11204-1-AP), PGAM1 (Proteintech, 16126-1-AP), HK2 (CST, 2867S), ALDOA (Proteintech, 11217-1-AP), TOMM22 (Proteintech, 66562-1-Ig), TOMM40 (Proteintech, 18409-1-AP), TOMM5 (Proteintech, 25607-1-AP), TOMM7 (ABclonal, A17711), TOMM20 (Santa Cruz, sc-17764), TOMM70 (Proteintech, 14528-1-AP*), SLC25A46* (Proteintech, 12277-1-AP), TID-1L/S (DNAJA3 – Santa Cruz, sc-18819), OXA1L (Proteintech, 21055-1-AP), NDUFA9 (Abcam, ab14713), TIMM23 (Santa Cruz, sc-514463), DNAJC19 (Life Technologies|AB /Invitrogen, 12096-1-AP), mtHSP70 (HSPA9 – Invitrogen, MA3-028), TIMM44 (Proteintech, 13859-1-AP), PAM16 (Proteintech, 15321-1-AP).

Secondary antibodies – IRDye® 800CW Goat anti-Rabbit IgG Secondary Antibody (LI-COR, 926-32211) and IRDye® 680RD Goat anti-Mouse IgG Secondary Antibody (LI-COR, 926-68070).

#### Confocal microscopy

Most confocal microscopy was performed using an Olympus FLUOVIEW FV3000 microscope in Galvano mode with a PlanApo N 60X 1.42 oil objective. The exception was for images appearing in panels Fig. 5C and D. For these, a Zeiss LSM 880 AiryScan Confocal Microscope was used. For live cell experiments, an okolab stage heater and CO_2_/air mixer kept samples at 37°C in a 5% CO2 environment. Live imaging was typically performed over the course of 1 – 2 hours. All confocal imaging of HeLa cells was performed in uncoated ibidi µ-Slide 8 Well chambered coverslips.

To measure PINK1-YFP intensity on mitochondrial with high and low MMP within the same cell, live HeLa^PINK1-YFP+MTS-mSc^ cells were loaded with MitoLite NIR following the manufacturer’s instructions for 15 – 20 minutes prior to imaging. Cells were imaged directly without additional washing. A midplane confocal image was obtained of each field and further processed in FIJI (Version 1.54m), using a custom FIJI macro script. Individual cells in the field were manually enclosed in an ROI using the polygon tool. The image was then duplicated and cropped around the cell of interest. The image was again duplicated, the background color was set to (0 0, 0), the images were downgraded to 8 bits, and the channels were split. MTS-mSc and MitoLite NIR masks were generated from duplicated images of the corresponding channels using the setThrehold (80, 255, “raw”) command followed by the run(“Convert to Mask”) command. A mitochondrial mask was generated from the union of the MTS-mSc and MitoLite NIR masks. A mask of high MMP mitochondria was generated by subtracting the MitoLite NIR mask from the mitochondrial mask. A mask of low MMP mitochondria was then generated by subtracting the high MMP mitochondria mask from the mitochondrial mask. Two groups of ROIs were then made from the high and low MMP mitochondria masks using the run(“Create Selection”) command to measure the intensity of each channel for each ROI. Measurements were made using the roiManager(“multi-measure measure_all”) command. The results table was exported as a .csv file. A custom Python script was used to collate data from the results .csv files for each cell. A similar process was used to measure mitochondrial PINK1-YFP from MitoLite NIR high cells following CTRL or TIMM23 KD, except that the MitoLite NIR mask was used to generate mito measurements and a non-mito mask was created by subtracting the MitoLite NIR mask from a cell mask defined by manual cell selection. For measurements of colocalization, cells were individually selected using the polygon tool, duplicated, and cropped as above. The “colocalization test” plug-in was then run, which measures the Pearson correlation coefficient for both the image and 20 control images produced by pixel randomization. The average Pearson coefficient from the random images (Rrand) was substrated from the observed Pearson coefficient (Robs) for each cell.

Immunocytochemistry was performed following fixation in 4% PFA n 1X PBS (diluted from paraformaldehyde 16% Aqueous Solution EM Grade, cat# 15700) for 10 minutes at room temperature. Cells were permeabilized by incubation with 0.25% Triton X100 for 10 minutes at room temperature. Cells were blocked for at least 15 minutes in 5% BSA in 1 X PBS. The primary antibody was diluted in 5% BSA in 1 X PBS at 1:500 ratio and incubated at room temperature for 1 – 2 hrs at room temperature or overnight at 4°C. Cells were washed three times in 1 X PBS. The secondary antibody (1:500) in 5% BSA 1 X PBS was added, and cells were incubated for 45 min at room temperature. The cells were washed three times in 1 X PBS and imaged in 1 X PBS.

#### Neuronal Staining

Neurons were cultured on 8 well glass bottom chamber slides (ibidi USA, Inc., 80827-90) and fixed in 4% PFA at d9, followed by permeabilization and block in 0.1% saponin (Sigma-Aldrich, S7900-100G) and 3% donkey serum (GeneTex, Inc., GTX73205) in PBS at room temperature for 30 minutes. Cells were incubated in primary antibodies prepped in permeabilization and block buffer overnight rocking gently at 4⁰C. Primary antibody was washed off with PBS and secondary antibodies were diluted (1:500) in permeabilization and block buffer and added to cells for 1hr rocking at RT. Secondary antibodies were washed off with PBS and DAPI (Invitrogen, 62248) was added 1:5000 in PBS for 5 minutes. Primary antibodies MAP2 (Proteintech, 17490-1-AP) and TUBB3 (BioLegend, 801201) both at 1:1000 dilution.

Secondary antibodies Goat anti-Rabbit IgG (H+L) Highly Cross-Adsorbed Secondary Antibody, Alexa Fluor™ Plus 555 (Invitrogen, A32732) and Goat anti-Mouse IgG (H+L) Highly Cross-Adsorbed Secondary Antibody, Alexa Fluor™ Plus 488 (Invitrogen, A32723) both at 1:500 dilution. Cells were imaged on an Olympus FLUOVIEW FV3000 confocal laser scanning microscope.

#### Clear Native Page and NAMOS assay

HeLa^PINK1-YFP^ cells were cultured with indicated sgRNA for at least 7 days before Clear Native Page sample collection. Samples were lysed and ran according to a previously published protocol ^59^. Briefly, cells were lysed in cold NativePAGE Sample Buffer (Invitrogen, BN2003) with 1% digitonin and 1X protease/phosphatase Inhibitor (CST, 5872S) and were placed on a rocker for 15 minutes at 4⁰C.

Samples were mixed by pipetting 10 times and then spun at 20,000g, 4⁰C, 30 minutes. Following BCA assay, 35 µg of cell lysate was incubated with 1µg of the indicated antibodies at room temperature for 1 hour (shaking). The whole volume of sample was loaded onto a NativePAGE Bis-Tris Mini Protein Gel 3- 12% (Invitrogen, BN-1001BOX) and were run in NativePAGE Running Buffer (Invitrogen, BN2001) supplemented with 0.05% Sodium Cholate (Sigma, C6445-10g) and 0.01% *n*-Heptyl-β-D-thioglucoside (Sigma, H3264) in the cold room (4⁰C). Gel was immediately imaged after running on an Amersham Typhoon 5 using 488nm cy2 laser-filter set combination. Total protein was measured via SimplyBlue SafeStain according to manufacturer’s protocol (ThermoFisher, LC6065) and imaged on a Cytiva Amersham ImageQuant 800.

Antibodies: TOMM22 (Proteintech, 66562-1-Ig), TOMM40 (Proteintech, 18409-1-AP), TOMM20 (Santa Cruz, sc-17764), and CHCHD2 (Proteintech, 66302-1-Ig).

#### Preparations that were submitted for Proteomics

##### Mitochondria Isolation and Insoluble/Soluble for Proteomics

Mitochondria were isolated from cells with CTRL or DNAJA3 sgRNA (4 replicates/per condition) according to a previously published protocol ^6,60^. Cells were homogenized with 40 strokes. Separation of insoluble and soluble fractions was completed according to a published protocol with some modifications ^40^. Briefly, extracted mitochondria were resuspended Triton X-100 buffer (20 mM Tris-HCl, pH 7.4, 150 mM NaCl, 2 mM EDTA, 1% Triton X-100), PI/PS (CST, 5872) and incubated for 30 minutes on ice (pipetting every 10 minutes). Cell lysates were centrifuged at 20,817g for 10 minutes at 4⁰C. Supernatant (soluble fraction) was moved to a new tube, aliquot was taken and added to 10% SDS to get a final solution of 5% SDS. Insoluble pellet was resuspended in Triton buffer, PI/PS, 5% SDS and agitated into solution. Samples were submitted for proteomics.

#### Whole Cell Lysate Preparation

Cells were pelleted in PBS at 4⁰C, 400g, 5 minutes, and resuspend in RIPA buffer (CST, 9806) and PI/PS (CST, 5872). The lysates were sonicated (QSonica Q800R3, Q800R3-110) for 10 minutes at 4⁰C followed by centrifugation at top speed (21,130g) 4⁰C, 5 minutes. BCA assay (Thermo Scientific, 23225) was performed on the supernatant. 40µg of lysate with 5% SDS was submitted for proteomics.

#### Immunoprecipitation of PINK1 YFP

Immunoprecipitation of HeLa^PINK1-YFP^ cells was preformed using GFP-Trap Magnetic Particles M-270 (chromotek, gtd) beads according to manufacturer’s instructions. Samples were eluted from beads in 30uL RIPA buffer (CST, 9806) + 5% SDS (Invitrogen, 15553-035) + PI/PS (CST, 5872) and the bound fraction was submitted for proteomics.

#### Mass spectrometry-based proteomics

Liquid chromatography-tandem mass spectrometry (LC-MS/MS) data acquisition was performed on an Orbitrap Ascend mass spectrometer (Thermo Fisher Scientific) coupled with a 3000 Ultimate high-pressure liquid chromatography instrument (Thermo Fisher Scientific) unless noted otherwise. Peptides were separated on an ES902 column (Thermo Fisher Scientific) with the mobile phase B (0.1% formic acid in acetonitrile) increasing from 3 to 18% over 61 min, then from 18% to 30% over 14 min. The LC MS/MS data were acquired in data-independent mode (DIA). Both MS1 and MS2 scans were performed in Orbitrap. For MS1 scan, the following parameters were used: mass range (m/z) = 380-985; resolution = 60K; the automatic gain control (AGC) = 4e5; RF lens = 60%. The parameters for MS2 scan were: mass range (m/z) = 145-1450; resolution = 15K; isolation window (m/z) = 10; window overlap (m/z) = 1; AGC = 1e5; maximum ion injection = 40 ms. The cycle time was set at 3 sec.

Spectronaut 19.5 software was used to analyze DIA data with the directDIA analysis method. Data were searched against Sprot Mouse database. Trypsin/P was set as specific enzyme with 2 max missed cleavages. Carbamidomethyl on cysteine was fixed modification. Acetyl (Protein N-tem and Oxidation on methionine were set as variable modifications. FDRs for PSM, peptide and protein level were 1%. MS2 abundances were used for protein quantitation. Cross-run normalization was performed. Unpaired t-test was used for hypothesis test.

Note: There were datasets acquired with data dependent method (DDA) as indicated in Table S10. The general method used for those samples was described previously (Shammas et al, 2022) with some main differences listed below. 1) Proteome Discoverer 3.1 (Thermo Fisher Scientific) was used for database search and quantitation analysis of DDA datasets. 2) ‘WCL exog vs endog PINK1’ and ‘mitochondria insoluble (I) / soluble (S) fractions’ datasets were acquired on an LC-MS system where an Orbitrap Ascend mass spectrometer (Thermo Fisher Scientific) was connected to a Vanquish NEO nano-HPLC (Thermo Fisher Scientific).

After quantification, proteomics data was further analyzed and visualized in Python, using custom scripts. Proteins were annotated for their mitochondrial localization and mitochondrial pathways using human MitoCarta3.0. Mitochondrial proteins with predicted strong presequences were identified using data from MTSviewer (https://mtsviewer.neurohub.ca/). Proteins were annotated as having a strong presequence if (1) they localized to mitochondria but not primarily the outer mitochondrial membrane, (2) their MTS1 score was >= 2, and (3) their MTS1 start position was < 20 amino acids from the N-terminus. For all proteomics experiments except AP-MS experiments, two-sided Student’s t-tests were performed using SciPy 1.14.0 Python library and values were corrected for multiple comparisons across all gene groups/features by calculating an FDR with the Benjamini–Hockberg procedure, using the statsmodels 0.14.1 Python library. Proteins were annotated as significant if they had an FDR < 0.05 and an absolute log_2_ fold-change of > 1 for most experiments; the exception was experiments in (Fig. 7G) where an absolute log_2_ fold-change of > 0.5 was used instead. For AP-MS experiment, statistics were performed in Perseus: missing data was first imputed using the “Replace missing values from normal distribution” function; a two-sided Student’s t-test was then performed within Perseus using default settings; finally, an FDR was calculated using the permutation method. For all proteomics data, volcano plots were generated using the Seaborn Python library. All other graphs were generated in Graphpad Prism 10.

#### Peroxidase staining and TEM of APEX-BFP expressing HeLa cells

HeLa^dCas9-BFP-ZIM3^ cells were transduced with dual RNA guides for the top eight proteins identified as PINK1-Parkin activators and expressing ER-BFP2-APEX targeted to the ER lumen. Previously cloned sgRNA plasmids were cut with restriction enzymes EcoRI and NHEI, followed by NEBuilder cloning with an APEX BFP gBlock. Cells transduced with sgRNAs were cultured on Aclar film coverslips (Electron Microscopy Sciences (EMS)) in 12-well plates. Coverslips were prepared by cutting Aclar into ∼15 mm squares that fit into wells of 12-well plates. Coverslips were rinsed in distilled water, then sterilized with 70% ethanol and transferred to a 12-well plate where they were rinsed ten times with autoclaved ultrapure water and rinsed once with culture medium before plating of cells. Cells were fixed 7 days after transduction for 30 minutes at room temperature in fixative containing 2% paraformaldehyde, 2% glutaraldehyde, and 50 mM calcium chloride in 0.1M sodium cacodylate buffer (EMS). Peroxidase staining was performed as previously described ^61^, with some modifications. After fixation, coverslips were rinsed twice in 0.1M sodium cacodylate buffer (cacodylate buffer), followed by a ten-minute incubation in cacodylate buffer containing 50 mM glycine (Sigma Aldrich) and a final rinse in cacodylate buffer. Coverslips were then incubated in 1-2 mL per well of cacodylate buffer containing 0.3 mg/mL of DAB (Sigma Aldrich) at room temperature in the dark. During DAB incubation, a 0.3% solution of hydrogen peroxide in cacodylate buffer was prepared by diluting 30% hydrogen peroxide (Sigma Aldrich) by 1:100 in cacodylate buffer. The resulting 0.3% hydrogen peroxide in cacodylate buffer was then added at a 1:100 dilution per well to the DAB containing cacodylate buffer in the 12-well plate, resulting in a final concentration per well of 0.003% hydrogen peroxide in 0.3 mg/mL DAB in cacodylate buffer, and the plate was returned to the dark. The intensity of the brown reaction product from the peroxidase reaction with DAB and peroxide was monitored by light microscopy and 10-12 minutes was determined to be the optimal development time (Fig. S2). Coverslips were rinsed twice with cacodylate buffer and then post-fixed with 3% glutaraldehyde in cacodylate buffer. For EM processing, coverslips were rinsed twice for 10 minutes in cacodylate buffer, and then stained with reduced osmium containing 1% osmium tetroxide (EMS) and 1% potassium ferrocyanide (Sigma-Aldrich) on ice for 60 minutes. Next, coverslips were rinsed three times with cacodylate buffer and dehydrated through a graded series of 5- minute ethanol rinses prior to infiltration in EmBED812 epoxy resin using the manufacturer’s hard resin formulation (EMS). To embed cells, Aclar coverslips were laid cell-side-up on a backing piece of Aclar, and then 2-3 gelatine capsules (EMS) filled with resin were inverted on top of the coverslips and placed in a 60°C oven for two days.

For sectioning, Aclar coverslips were peeled away from the resin and en face ultrathin sections of embedded cells were cut to a thickness of 70 nm using an ultramicrotome (EM UC7, Leica Microsystems) with a diamond knife (DiATOME). Sections were picked up on 1 mm-slot copper grids with a thick formvar support film (EMS), and post-stained with 3% Reynold’s lead citrate (EMS) for 1 minute.

Sections were viewed using a JEOL1400 Flash transmission electron microscope (JEOL USA, Inc.) operated at 120 KV. Images were recorded with a Biosprint29 CMOS detector (Advanced Microscopy Techniques). Images were viewed and adjusted for display using Adobe Photoshop Software version 2023 (Adobe). Linear adjustments to black and white points were made using the Levels tool to optimize contrast. Figures were prepared using Adobe Illustrator software version 2023 (Adobe).

#### Assessment of TEM ultrastructure of mitochondria transduced with sgRNAs

Thin sections of cells expressing sgRNAs were searched and cells with ER that was darkly stained with the APEX-DAB reaction product indicating unequivocally high transduction level were recorded for analysis. At least ten cells from of each transduction condition were analyzed. First the ultrastructure of mitochondria in control cells were surveyed to establish the baseline appearance of mitochondria in control HeLa cells. Then cells that were transduced with sgRNA to knockdown mitochondrial proteins were reviewed and scored as exhibiting normal versus abnormal mitochondria. Then the unusual features in mitochondrial cristae structure, matrix, and mitochondrial shape in cells deemed abnormal were recorded and are presented in Table S2.

#### Correlative light and electron microscopy

Transduced HeLa cells were plated on either 15 mm Alcar film coverslips (EMS) or glass coverslips (18 mm or 25 mm diameter) (Fisher Scientific). For CLEM, prior to sterilization and cell plating, coverslips were shadowed with a 4 nm-thick layer of carbon deposited using a vacuum evaporator system (Leica Microsystems). The coverslips were cleaned with water and ethanol and air dried. Then a 10 mm-wide finder grid (EMS) was placed on the center of a coverslip before loading it in the carbon evaporator system. The layer of carbon was deposited on the surface of the coverslip, and presence of the finder grid created numerical grid pattern that was visible by light microscopy. After 7 days of transduction, cells were fixed with 4% paraformaldehyde in 0.1M sodium cacodylate buffer (cacodylate buffer) (EMS) and then either loaded into a cell chamber in cacodylate buffer (ThermoFisher Scientific) or rinsed and mounted on a glass slide (Fisher Scientific) in ProLong Gold Antifade mountant (ThermoFisher Scientific) for confocal imaging.

Confocal imaging was performed with either an Olympus FLUOVIEW FV3000 Microscope (Fig 3D and Fig S5S) or Zeiss LSM 880 AiryScan Confocal Microscope l (Fig. 5D) and images are shown as z-projections. Transmitted light images were taken to record the location of the cells of interest relative to the carbon shadowed grid. Following confocal imaging, coverslips in mounting medium were released from glass slides by soaking the slide in cacodylate buffer until the coverslip was free. Coverslips were rinsed in cacodylate buffer and then fixed in 2% glutaraldehyde (EMS).

EM processing was carried out as for APEX-BFP expressing HeLa cells, with the added step of staining with 1% uranyl acetate (EMS) in acetate buffer, pH 5.2 (EMS) overnight at 4°C. Specifically, after reduced osmium staining, cells were rinsed with cacodylate buffer and then 50 mM acetate buffer pH5.2 (EMS) before overnight incubation in uranyl acetate. The next day, cells were rinsed in acetate buffer before dehydration and embedding. To embed correlative samples, after infiltration with resin, a resin-filled gelatin capsule was inverted and placed directly over the location on the correlative grid pattern where the cell of interest was located. Glass coverslips were laid cell-side-up on a glass slide prior to placement of the inverted gelatine capsule, and Aclar coverslips were laid on a piece of Aclar film before placement of the gelatine capsule and polymerized in a 60°C oven for two days. Glass coverslips were separated from the resin by heating the back of the glass slide with a flame until the resin block separated from the glass; Aclar coverslips were peeled away from the resin. The carbon grid transferred to the surface of the resin block was used to locate the cell of interest for thin sectioning. Serial sections were cut at 70 - 140 nm thickness and picked up on 1-mm slot grids (EMS), post stained with 3% Reynold’s lead citrate (EMS) before viewing on the electron microscope as described above. Alignment of the confocal and EM images was achieved using the TrakEM2 plugin for Fiji ^62^.

#### TOMM5 Rescue

TOMM5 WT or TOMM5 1-39 (truncation after residue 39) were cloned into the doxycycline inducible pSBtet-RN a gift from Eric Kowarz (Addgene plasmid # 60503 ; http://n2t.net/addgene:60503 ; RRID:Addgene_60503) ^63^. Briefly, pSBtet-RN was digested with SfiI and ligated via NEBuilder with either HA-TOMM5 WT or HA-TOMM5 1-39 gene blocks. Ligations were transformed in One Shot TOP10 Chemically Competent *E. coli* (Invitrogen, C404003) according to manufacturer’s instructions. Single colonies were picked and grown up in LB broth, mini prepped with the QIAprep Spin Miniprep Kit, and sequenced (sequencing primer - AGCTCGTTTAGTGAACCGTCAGATC). These plasmids were transfected via FuGENE HD according to manufacturer’s instructions into HeLa PINK1 KO PINK-YFP dCas9-BFP-ZIM3 cells with the transposase (pCMV(CAT)T7-SB100 was a gift from Zsuzsanna Izsvak (Addgene plasmid # 34879) ^64^. Briefly, a ratio of 3:1 FuGENE HD was used (3µL:1µg) and for DNA a 1:20 ratio transposase:plasmid. Following FACS sorting to obtain a polyclonal population the cells were transduced with CTRL or TOMM5 sgRNA following the above sgRNA transduction protocol. The cells were maintained in +/- 1 µg/mL doxycycline for the duration of the experiment. Cells were treated with vehicle or 10 µM CCCP for 4 hours and run on the CellStream after at least 7 days post sgRNA transduction.

#### Statistical analysis

All statistics were calculated in Graphpad Prism 10, except as noted above for the proteomics experiments. When the variance was similar among the groups and the data was normally distributed, standard one-way ANOVA was performed when comparing differences between more than 2 groups, and an unpaired, 2- tailed t-test was performed when comparing 2 groups. Where these assumptions were not true, multiple groups were compared using a Brown-Forsythe and Welch ANOVA was performed followed by Dunnett’s T3 multiple comparisons test, and for comparison to two groups a Mann-Whitney test was performed.

---

**Supplemental Figure 1.**
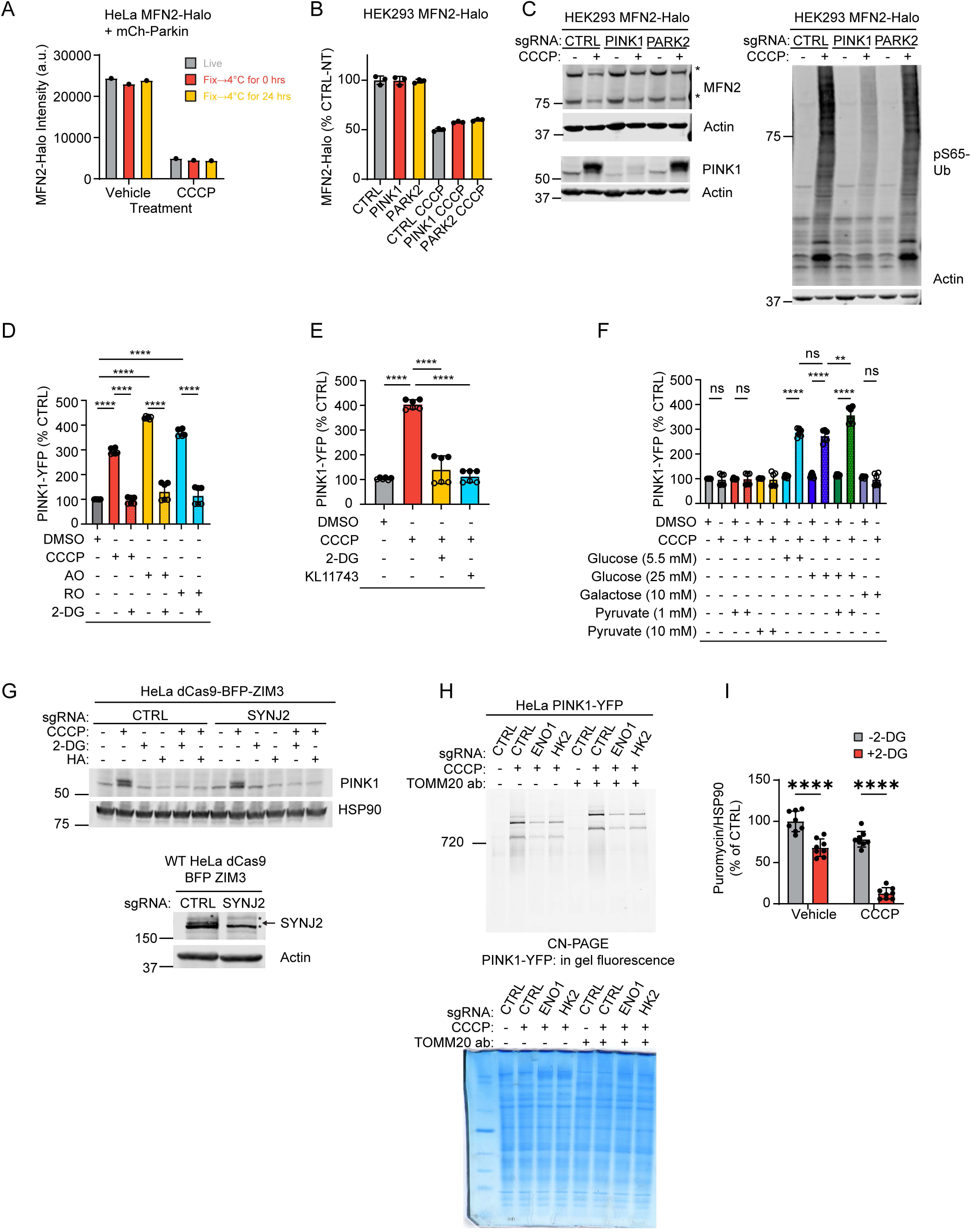
MFN2-HaLo reporter validation and glycolysis is required for PINK1-YFP stabilization following OXPHOS impairment. (A) Flow cytometry of HeLa^MFN2-Halo+mCh-Parkin^ cells +/- 10 µM CCCP 4hrs with or without fixation. N = 1 independent experiment. (B) Flow cytometry of HEK293^MFN2-Halo^ cells +/- 10 µM CCCP O/N. N = 3 replicates on 1 occasion. (C) Representative immunoblots of endogenously tagged and untagged MFN2 alleles (top and bottom band, respectively) +/- 10 µM CCCP 4 hrs from HEK293^MFN2-Halo^ cells. * denotes ubiquitinated MFN2 band. N = 2 replicates on 1 occasion. (D) Flow cytometry measurements in HeLa^PINK-YFP^ cells treated with 10 µM CCCP, and 8 µg/mL antimycin + 10 µg/mL oligomycin, 1 µM rotenone + 10 µg/mL oligomycin, and/or 10 mM 2-DG, for 4 hrs. **** p ≤ 0.0001 Error bars mean +/- SD. N = 6 independent experiments from 2 separate transductions. (E) Flow cytometry measurements in HeLa^PINK1-YFP^ cells treated with 10 µM CCCP, 10 µM KL11743 and/or 10 mM 2-DG, for 4 hrs. **** p ≤ 0.0001 Error bars mean +/- SD. N = 6 independent experiments from 2 separate transductions. (F) Flow cytometry measurements in HeLa^PINK1-YFP^ cells treated with 10 µM CCCP +/- in glucose and pyruvate-free DMEM supplemented with glucose, galactose, and/or pyruvate at the indicated concentrations for 4 hrs. **** p ≤ 0.0001 Error bars mean +/- SD. N = 6 independent experiments from 2 separate transductions. (G) Representative immunoblots of HeLa^dCas9-BFP-ZIM3^ cells treated with 10 µM CCCP, 10 µM HA, and/or 10 mM 2-DG 4 hrs. * denotes non-specific bands, arrow denotes position of SYNJ2 band. N = 2 independent experiments. (H) CN-PAGE separated PINK1-YFP complexes visualized by in gel fluorescence as in (Fig. 5J) (top) and total protein measured via SimplyBlue SafeStain (bottom). HeLa cells were cultured with indicated sgRNA for at least 7 days before CN-PAGE sample collection. N = 3 replicates from at least 2 transductions. (I) Quantification of immunoblots represented in figure 2G. **** = p ≤ 0.0001. Error bars mean +/- SD. N = 3 independent experiments.

**Supplemental Figure 2.**
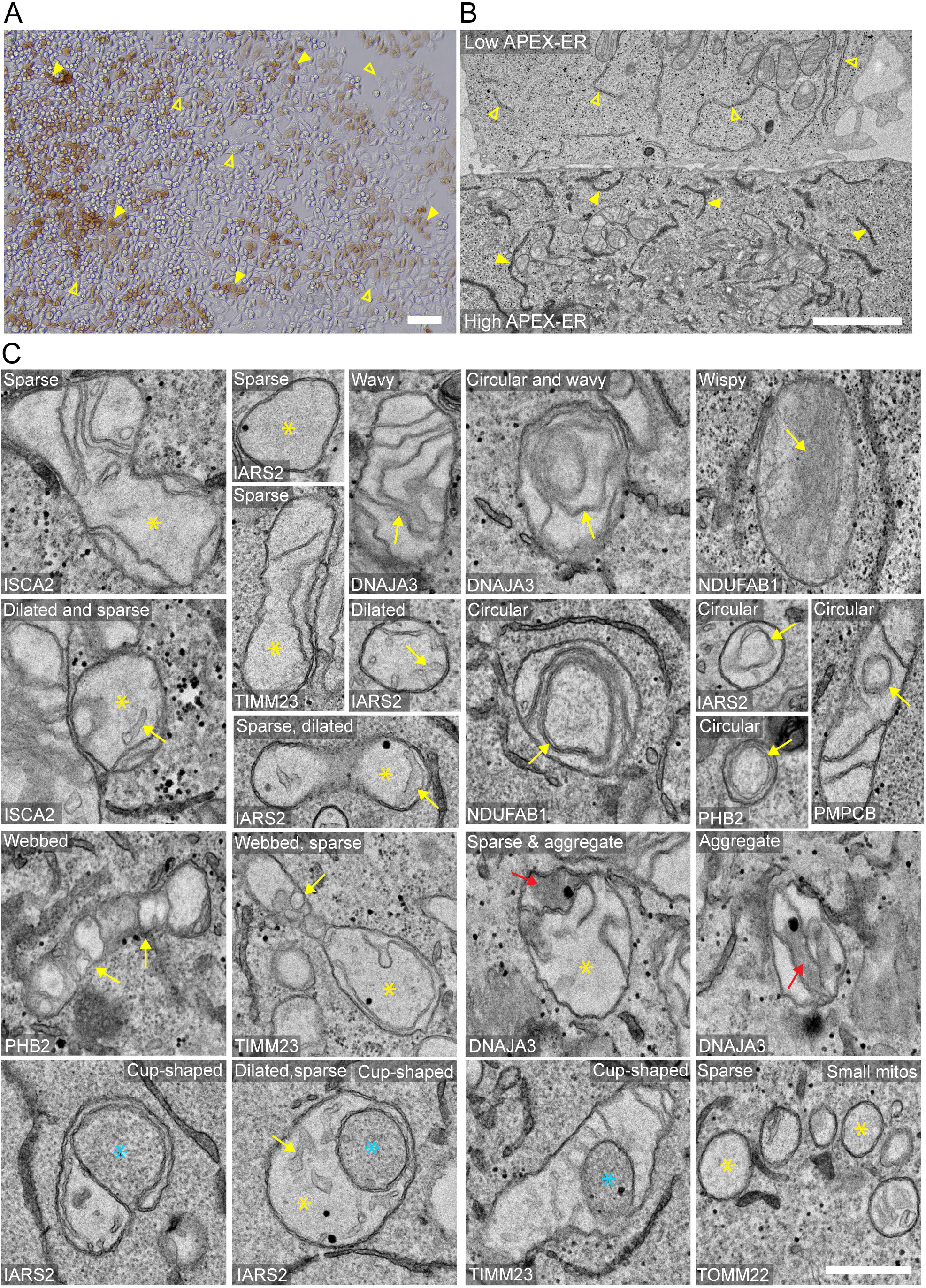
APEX-ER staining reliably identifies HeLa cells with knockdown of mitochondrial proteins for assessment of mitochondrial ultrastructure by TEM. (A) Transmitted light image of HeLa^dCas9-BFP-ZIM3^ cells shows the brown DAB reaction product after ∼12 minutes of development time in cells transduced with APEX-ER (solid arrowheads) compared to cells expressing low or no APEX-ER (open arrowheads). Scale bar = 100 µm. (B) TEM image of a cell with low or no APEX-ER staining in the ER lumen (open arrowheads) next to a cell with darkly stained ER lumen (closed arrowheads) indicating a high level of APEX-ER expression. Scale bar = 2 µm. (C) Examples of abnormal ultrastructural features of mitochondria expressing sgRNAs for the protein indicated in the lower left corner of each image. Yellow arrows indicate the abnormal cristae feature listed at the top of each image; yellow asterisks indicate areas with sparse cristae. Red arrows indicate the fluffy aggregate observed in the matrix of cells transduced with sgRNA DNAJA3. Blue asterisks indicate cytosol enclosed cup-shaped mitochondria. In some cases, examples shown were cropped from the same cell. Scale bar = 500 nm.

**Supplemental Figure 3.**
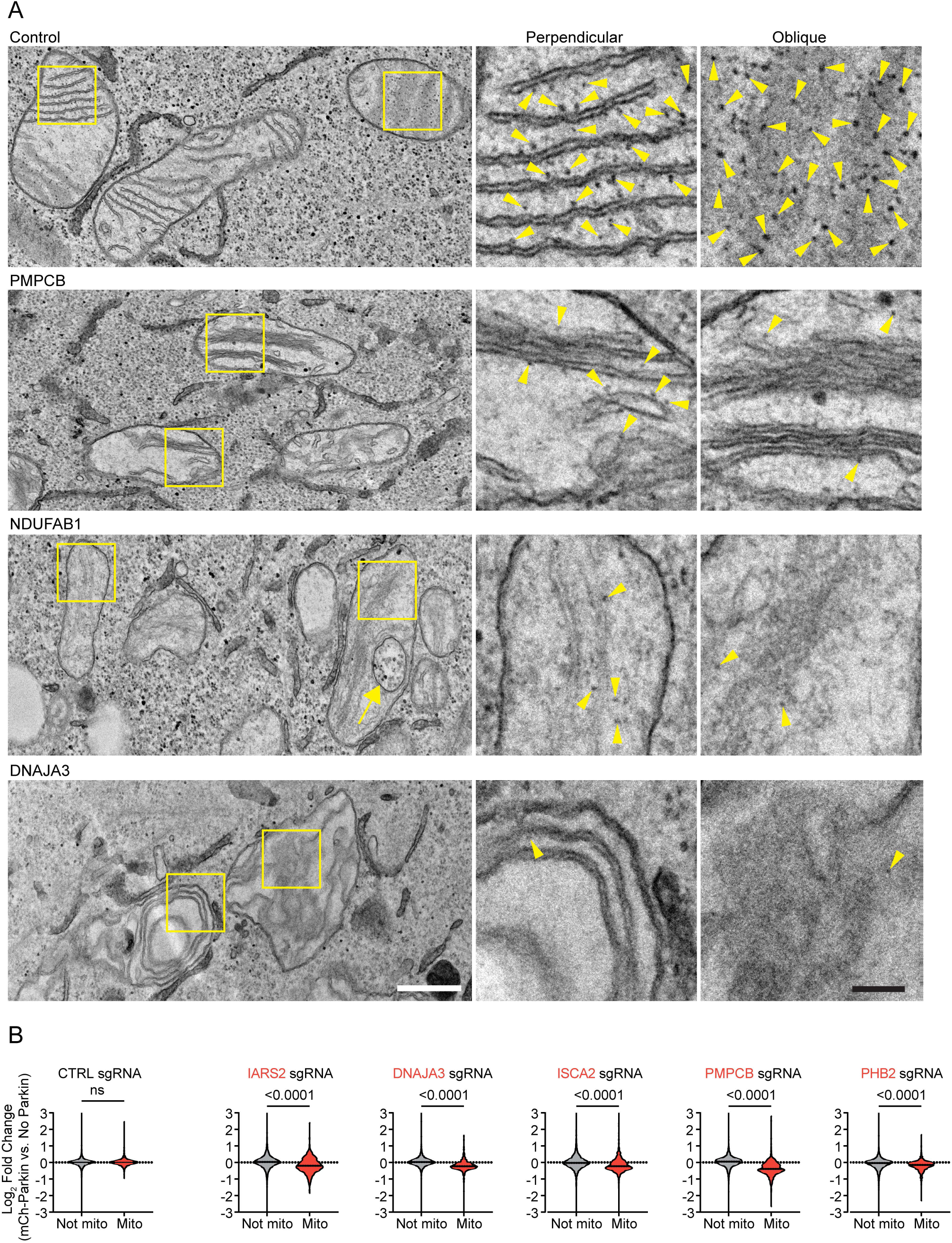
Cristae membranes may lose F_1_F_O_-ATP synthase by TEM following mitochondrial perturbations that activate PINK1-Parkin. (A) EM shows loss of cristae-associated punctate densities that may correspond to the F_1_F_O_-ATP synthase in HeLa^dCas9-BFP-ZIM3^ cells transduced with sgRNA targeting PMPCB, NDUFAB1, and DNAJA3. In control cells (top), numerous punctate structures decorate cristae. Yellow boxed areas shown to the right with yellow arrowheads indicate the abundance of presumed F_1_F_O_-ATPase synthase visible along cristae cut perpendicularly in thin sections and when cristae are sectioned in an oblique orientation. Possible F_1_F_O_-ATP synthase particles are observed along cristae in mitochondria of cells expressing sgRNA for PMPCB and NDUFAB1, but at a reduced frequency. Possible F_1_F_O_-ATP synthase densities are virtually absent along cristae of mitochondria in cells transduced with sgRNA for DNAJA3. White scale bar = 500 nm; black scale bar = 100 nm. (B) LFQ performed as described in (Fig. 3H) comparing knockdowns in HeLa cells with mCh-Parkin vs. HeLa cells with no Parkin. HeLa cells without Parkin are from the same data that appears in (Fig. 3H).

**Supplemental Figure 4.**
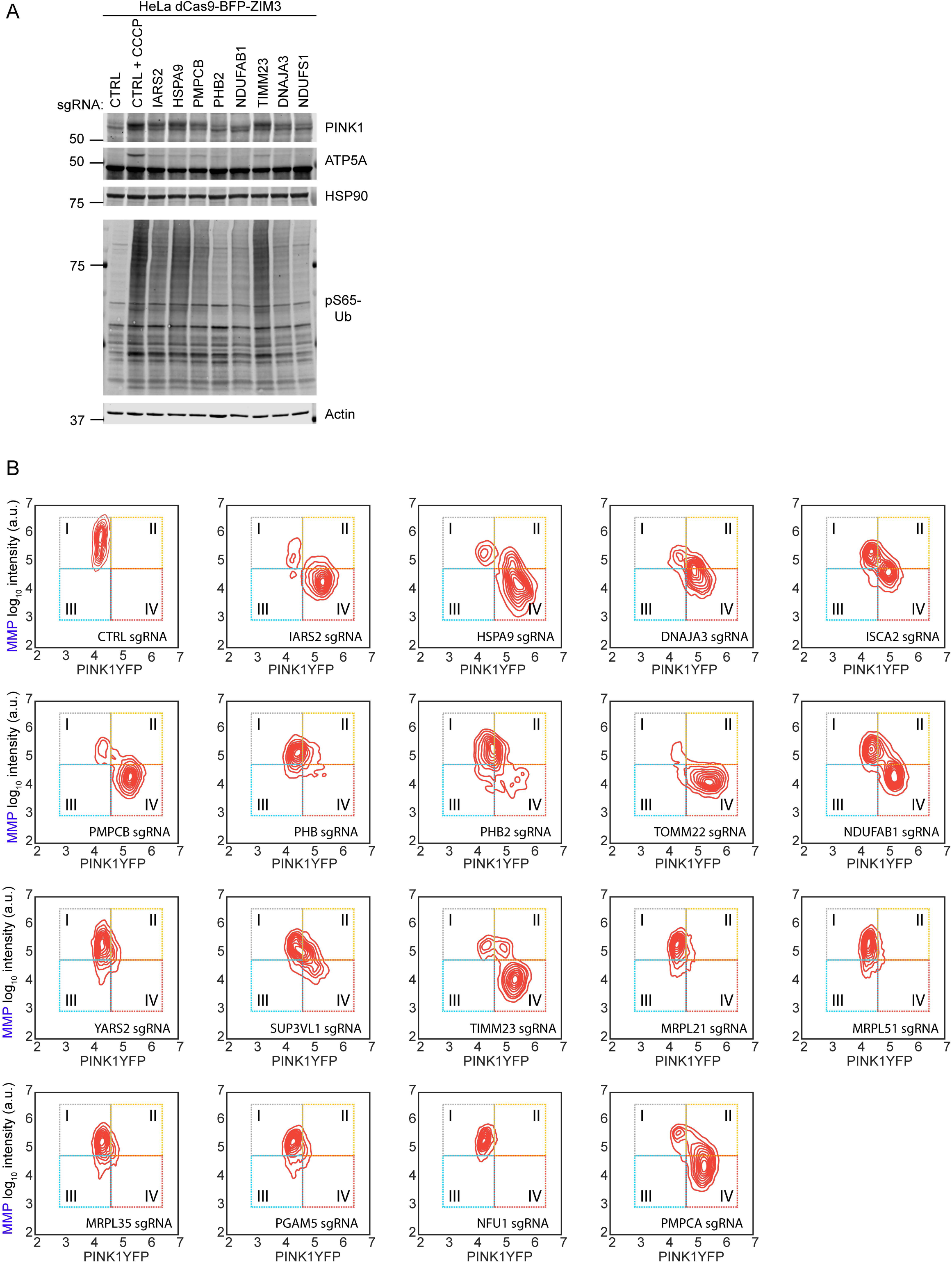
Top PINK1-Parkin activators stabilize endogenous PINK1 and lower MMP. (A) Representative immunoblots of HeLa^dCas9-BFP-ZIM3^ cells treated with 10 µM CCCP for 4 hrs or transduced with indicated sgRNAs. N = 3 independent experiments. (B) Representative 2D kernel density plots comparing single-cell PINK1-YFP intensity and intensity of the MMP sensitive dye MitoLite NIR as in (Fig. 4F).

**Supplemental Figure 5.**
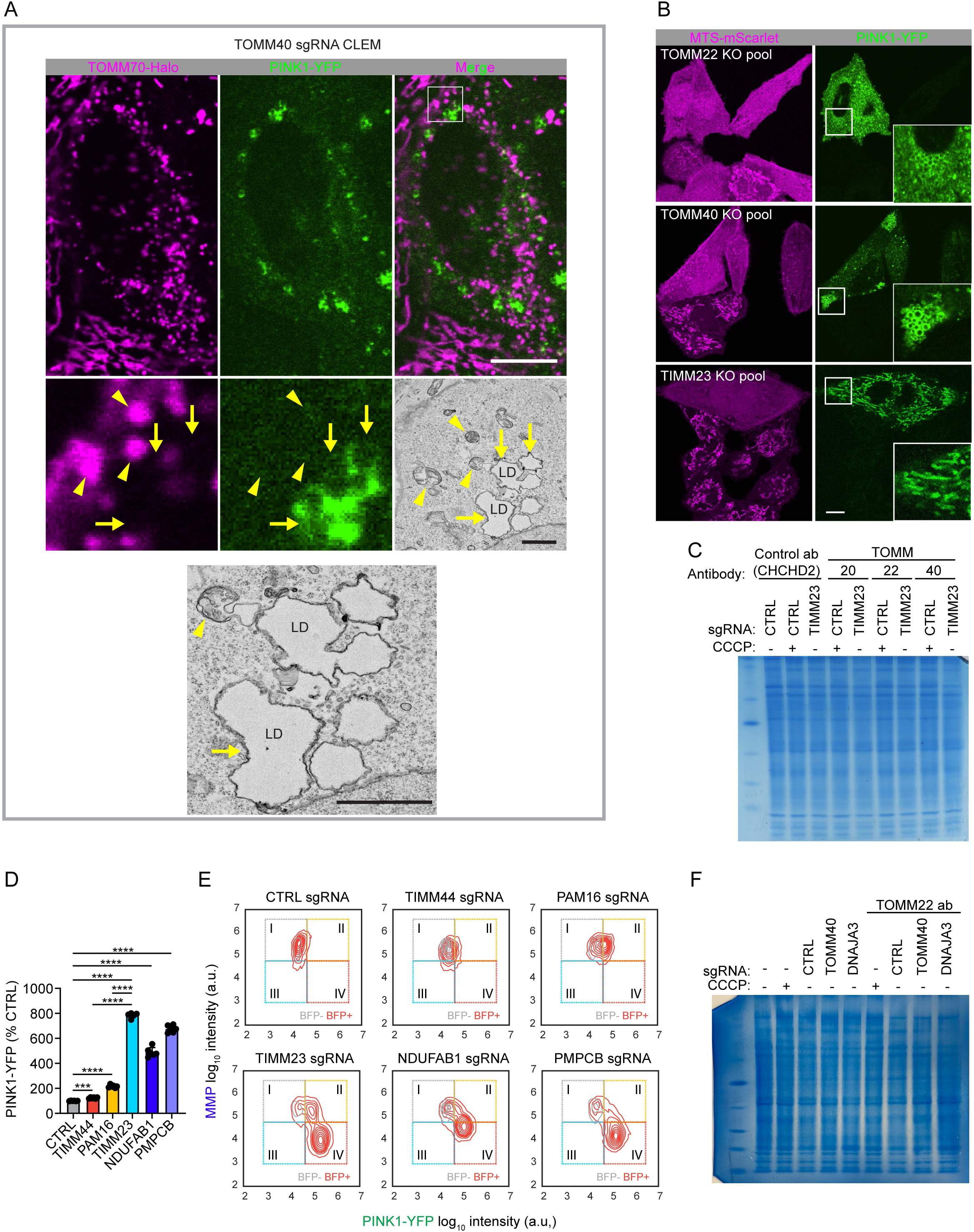
PINK1 is differentially affected by loss of the TOM complex, TIM23, and the PAM import motor. (A) CLEM of HeLa^PINK1-YFP^ cells endogenously tagged with TOMM70-Halo and transduced with sgRNA targeting TOMM40. Fluorescence confocal microscopy shows that in cells with sgRNA-mediated knockdown of TOMM40, the mitochondrial marker, TOMM70-Halo and PINK1-YFP were not colocalized. The same cell examined at the EM level and aligned with the confocal image showed TOMM70-Halo label colocalized with small round mitochondria while the PINK1-YFP signal localized to a cluster of lipid droplets (LD). Yellow arrowheads indicate Halo-labeled mitochondria. Yellow arrows indicate site of PINK1-YFP accumulation around lipid droplets. White scale bars = 10 µm, black scale bars = 1 µm. (B) Representative confocal images of TOMM22, TOMM40, TIMM23 KO pools in HeLa^PINK1-YFP+MTS-mSc^ cells showing PINK1-YFP accumulates in the same pattern as observed by CRISPRi. Images were obtained 7 or 8 days after electroporation with Cas9 ribonucleoprotein complexes. Scale bar 10 = µm. (C) Total protein measured via SimplyBlue SafeStain of same gel as in (Fig. 5J), demonstrating equal loading. (D) Flow cytometry of HeLa^PINK1-YFP^ cells. ***≤ 0.001, **** p ≤ 0.0001 Error bars mean +/- SD. N = 6 replicates from 2 independent transductions. (E) Representative 2D kernel density plots comparing single-cell PINK1-YFP intensity and intensity of the MMP sensitive dye MitoLite NIR as in (Fig. 4F). Total protein measured via SimplyBlue SafeStain of same gel in (Fig. 7D), demonstrating equal loading.

